# DefensePredictor: A Machine Learning Model to Discover Novel Prokaryotic Immune Systems

**DOI:** 10.1101/2025.01.08.631726

**Authors:** Peter C. DeWeirdt, Emily M. Mahoney, Michael T. Laub

## Abstract

Anti-phage defense systems protect bacteria from viruses. Studying defense systems has begun to reveal the evolutionary roots of eukaryotic innate immunity and produced important biotechnologies such as CRISPR-Cas9. Dozens of new systems have been discovered by looking for systems that co-localize in genomes, but this approach cannot identify systems outside defense islands. Here, we present DefensePredictor, a machine-learning model that leverages embeddings from a protein language model to classify proteins as defensive. We applied DefensePredictor to 69 diverse *E. coli* strains and validated 45 previously unknown systems, with >750 additional unique proteins receiving high confidence predictions. Our model, provided as open-source software, will help comprehensively map the anti-phage defense landscape of bacteria, further reveal connections between prokaryotic and eukaryotic immunity, and accelerate biotechnology development.

## Introduction

Bacteria face a relentless battle with viruses known as bacteriophages. In some environments, these phages can drive the turnover of 10-25% of all bacteria on a daily basis (*1*). The intense selective pressure to evade or survive infection has driven the evolution of numerous anti-phage defense systems, including, most famously, restriction-modification and CRISPR-Cas systems. Owing to their molecular specificity, these systems have been powerfully repurposed for genetic and genome engineering. Identifying new anti-phage defense systems may yield the next generation of precision molecular tools, while also shedding important light on the ongoing arms race between bacteria and phages. Notably, recent work has also indicated that many components of the mammalian innate immune system are homologous to, and likely originate from, bacterial proteins that function in anti-phage defense (*2*, *3*).

The full complement of bacterial anti-phage defense systems remains unknown. Several systematic searches for new defense systems leveraged the tendency of some defense genes to co-localize in genomes by looking for uncharacterized genes in these so-called ‘defense islands’ (*4*). These guilt-by-association methods have identified and validated 59 defense systems (*5*–*7*). A complementary experimental approach involving the screening of a library of genomic fragments from diverse *Escherichia coli* genomes identified 21 new defense systems, most of which reside within mobile genetic elements not defense islands (*8*). Other studies have bioinformatically mined mobile elements to identify yet more systems (*9*–*12*).

Despite their successes, these prior approaches have fundamental limitations. The guilt-by-association approach focuses exclusively on systems near known defense genes, and not all, and perhaps a minority of, systems are located in defense islands (*8*). The experimental screening approach is laborious and not done to saturation. Thus, we likely have a vast underestimate of the full landscape of anti-phage defense in bacteria, and lack the tools to systematically identify systems with high speed, sensitivity, and specificity.

Here, we built a machine learning model called DefensePredictor that can predict defense systems. Our model is a gradient boosting classifier built on top of embeddings from a protein language model (PLM). PLMs learn statistical relationships between amino acids within a protein by predicting the identity of masked amino acids from the surrounding protein context, analogous to the distributional hypothesis in linguistics in which the meaning of a word can be learned by the context in which it occurs (*13*, *14*). DefensePredictor outperforms guilt-by-association models on a held-out test set of defense genes. We used our model to identify novel defense genes in diverse strains of *E. coli* with an experimental validation rate of 42%, yielding 45 new defense systems, although the true positive rate of the model is likely higher as some systems may defend against phages that were not tested. Some new systems feature domains, *e.g.* nucleases, found in previous defense systems but in different contexts or configurations. Seven new systems contain domains not previously implicated in defense and we show each is essential for protection. These new domains include a metallophosphatase homologous to the human innate immune gene *SMPDL3A*. Collectively, our findings demonstrate that DefensePredictor represents a robust and powerful pipeline for identifying anti-phage defense systems. We provide DefensePredictor as an open-source Python package, enabling the rapid prediction of defense systems in any prokaryotic genome in under five minutes.

## Results

### DefensePredictor achieves high precision and recall on held out test data

To build a machine learning model to predict novel defense genes, we first searched ∼17,000 assembled genomes, representative of the taxonomic diversity of prokaryotes (Fig. 1A), for known defense systems using DefenseFinder (*10*). We identified ∼244,000 homologs that belong to complete defense systems. These genes represented our positive set for machine learning. To obtain our negative gene set, we identified ∼14 million genes with Gene Ontology (GO) or Clusters of Orthologous Genes (COG) annotations typically associated with non-defensive functions, *e.g.*, translation, membrane transport, etc. To reduce redundancy in both the positive and negative gene sets, we clustered proteins at 80% coverage and 30% identity using MMseqs2 (*15*), and randomly selected a maximum of five proteins per cluster.

**Figure 1.**
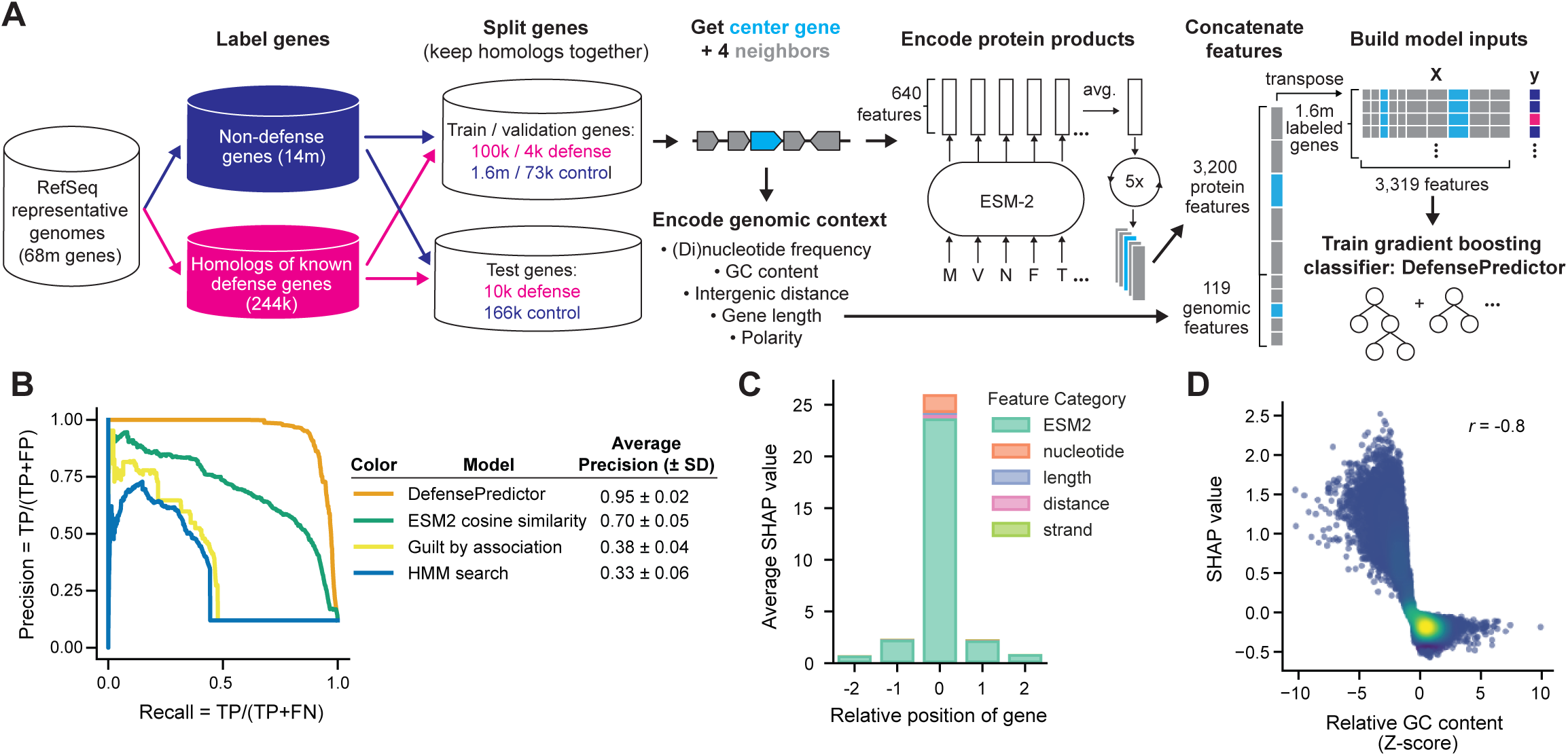
DefensePredictor shows strong performance on held out test data. (**A**) Pipeline for building DefensePredictor. Dark blue elements represent control genes, pink elements represent known defense genes. Light blue elements represent center genes, grey elements represent neighboring genes. m, million; k, thousand. (**B**) Precision-recall curves on held out test data. TP, True Positive; FP, False Positive; FN, False Negative. (**C**) SHAP values summed for each true positive gene (n= 8,653) and feature category and averaged across all true positive genes in the test set. Position 0 represents the center gene; other positions are neighboring genes. (**D**) SHAP value of GC content versus z-scored GC content for all center genes in the test set. Color represents density, with yellow being the densest area of the plot. Pearson correlation is indicated.

To split the defense genes into training, validation, and test sets, we first clustered the 1,018 HMM profiles from DefenseFinder. We connected two profiles if they both aligned to one or more of the same protein(s) in the set of representative genomes, or if they were defined as exchangeable by DefenseFinder (e.g. the CBASS transmembrane effector is exchangeable with the CBASS phospholipase effector). In our splits, we kept together all proteins that aligned to an HMM of the same cluster, resulting in a training set of ∼100,000 defense genes from 229 HMM clusters, a validation set of ∼4,000 defense genes from 14 HMM clusters, and a test set of ∼10,000 defense genes from 40 HMM clusters (table S1). By splitting the defense genes in this way, we can evaluate whether a trained model can identify defense genes that are not simply homologs or exchangeable with genes from known systems. We split our negative genes in a similar manner, keeping together proteins from the same MMseqs2 protein cluster. This resulted in a training set with ∼1.6 million negative genes from ∼940,000 clusters, a validation set with ∼73,000 negative genes from ∼42,000 clusters, and a test set with ∼166,000 negative genes from ∼96,000 clusters.

To create a feature matrix for training, we took each gene in the training set, hereafter called the ‘center gene’, and the two neighboring genes on either side. For each center and neighboring gene we built a representation of its protein product using the PLM Evolutionary Scale Model 2 (ESM2). ESM2 generates a 640-dimensional embedding for each amino acid in a protein, which we averaged across each coding sequence to obtain our protein-level representations (*14*, *16*). We concatenated the average embeddings for each center gene and its four gene neighbors to obtain a 3,200-dimensional vector of protein features.

To also capture genomic context information, for each center gene and its gene neighbors we calculated nucleotide and dinucleotide frequencies, GC content relative to the genome, gene length, the distance between each pair of consecutive genes, and orientation of the neighboring genes relative to the center gene, yielding 119 genomic features total. We concatenated the genomic and protein features to obtain a final feature vector size of 3,319. We calculated these features for all ∼1.6 million positive and negative genes in our training set to obtain our training feature matrix. Similarly, we built a feature matrix for the genes in our validation set, which we used to tune model hyperparameters (see Methods). Using the training feature matrix, we fit a gradient boosting classifier with the selected hyperparameters. This classifier combines predictions from thousands of decision trees, each of which partitions genes based on their feature values and estimates a probability of defense for each partition (*17*). We call this model DefensePredictor.

To evaluate the performance of DefensePredictor, we first used the model to predict the probability of defense for each gene in the held-out test set. Using the true labels for these genes, we calculated the precision (the proportion of predicted defense genes that are actually defensive) and recall (the proportion of actual defense genes that are predicted) of the model over descending probability thresholds (Fig. 1B). As a summary statistic, we calculated the Average Precision (AP) across all recall cutoffs. We found that DefensePredictor achieved an AP of 0.95, indicating that our model recovers unseen defense genes with high precision and recall.

To evaluate DefensePredictor’s performance on different families of defense proteins, we examined the model’s predictions for each of the 40 HMM clusters in the positive test set of genes. For 30 of 40 clusters, more than 50% of member proteins were predicted as defensive using a probability cutoff of 0.5 (fig. S1A). Thus, DefensePredictor can identify a wide range of defense protein families, but has a false negative rate of ∼25% at the protein family level.

We next compared DefensePredictor to a guilt-by-association method in which we assessed, for each protein cluster containing at least one member in the test set (positive or negative), whether the proteins in that cluster were frequently encoded in genomes near known defense genes (see Methods). We ranked all test genes based on the significance of their cluster’s association (measured using Fisher’s exact test) and calculated the precision and recall at descending significance thresholds. Using this method, we obtained an AP of 0.38, maintaining a high precision (∼0.75) until a recall of ∼0.25, at which point performance dropped.

We also assessed how well remote homology detection techniques recovered defense genes in the test set. To do so, we searched the proteins in the test set with the HMMs that were used to identify defense genes in the training set. We then ranked test genes based on the Expect (E) value of their most significant alignment. Calculating the precision and recall at increasing E-value cutoffs yielded an AP of 0.33. As a second benchmark of remote homology detection, we calculated the cosine similarity between the average ESM2 embedding of each test protein and the average ESM2 embedding of each defense protein in the training set, an approach shown to enable remote homology detection (*18*). We ranked test genes based on their maximum cosine similarity and calculated precision and recall at decreasing levels of similarity, obtaining an AP of 0.7. Taken together, our analyses indicate that DefensePredictor outperforms guilt-by-association and remote homology detection approaches and can identify truly novel defense genes.

Next, we aimed to understand the individual features that DefensePredictor uses to identify systems. To do so, we calculated SHapley Additive exPlanation (SHAP) values, which estimate how much each input feature contributes to a model’s predictions (*19*). The four highest average SHAP values came from individual ESM2 features for the center gene (Fig. 1C, table S2). The fifth most important feature was (low) G/C content of the center gene relative to the host genome (Fig. 1D). Known defense genes tend to have lower G/C content than their host genomes (*20*, *21*) (fig. S1B). Thus, this result suggests that DefensePredictor captured fundamental biological patterns to make its predictions. Nevertheless, ESM2 features of the center gene were the most important contributor to predictions (average SHAP value = 23.7).

SHAP values can also be summed for individual genes and used to assess how neighboring genes contributed to DefensePredictor’s outputs. We calculated these summed SHAP values and inspected the annotations of the neighboring genes that contributed most positively to predictions (table S2). We noted an enrichment of neighbor genes annotated as anti-phage defensive or encoding transposases or integrases (fig. S1C). These results suggest that DefensePredictor learned to leverage the clustering of defense genes and the association between defense genes and mobile elements to make its predictions.

### Prediction of novel defense genes in a diverse collection of *E. coli* strains

Next, we used DefensePredictor to identify systems in a set of 69 diverse *E. coli* strains, primarily from the ECOR collection (*22*), which captures much of the pangenome diversity of this species (table S3). Based on experimental validation studies (see below), to define a gene as defensive in our *in-silico* analyses we used a stringent probability cutoff of 0.999, or equivalently a log-odds cutoff of 7.2, where the log-odds of a gene is defined as 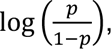 with *p* being its predicted probability of defense. Using this cutoff, the model predicted that ∼2,500 out of ∼321,000 genes from these strains encode defense proteins (Fig. 2A). In contrast, DefenseFinder identified only 395 proteins that were part of a complete system across all strains. Both approaches agreed on a core set of 325 proteins (Fig. 2B), 308 of which were represented in the training set, so this strong overlap is unsurprising.

**Figure 2.**
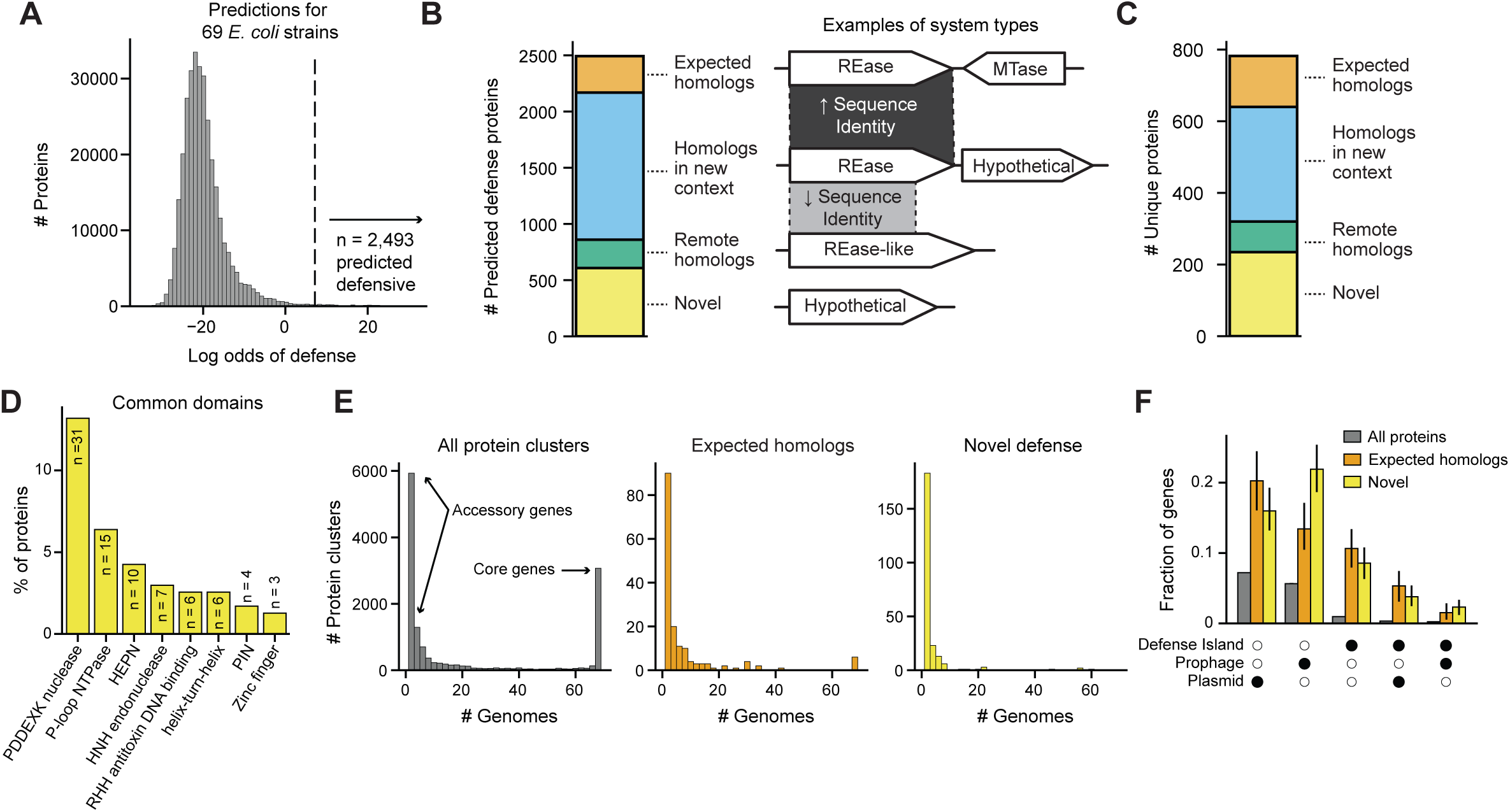
DefensePredictor identifies hundreds of predicted defense genes in 69 diverse *E. coli* strains. (**A**) Histogram of log-odds for 321,347 genes from 69 strains. Vertical line at a log-odds of 7.2 indicates a strict threshold for labeling a gene predicted defensive. (**B**) Left: number of proteins belonging to one of four categories of predicted defense genes. Right: examples of the types of predicted defense genes. Shaded areas represent regions of homology between genes, with darker regions representing stronger sequence similarity. REase, restriction endonuclease; MTase, methyltransferase. (**C**) Same as (**B**), after selecting one unique protein per homologous cluster. (**D**) Frequency of domain families found in unique predicted novel defense proteins. RHH, ribbon-helix-helix; HTH, helix-turn-helix. (**E**) Histogram of the number of genomes each protein cluster is encoded by. Protein clusters encoded by five or fewer genomes are considered part of the accessory pangenome and clusters encoded by all genomes are considered part of the core pangenome. (**F**) Frequency of genes residing in defense islands (`±`10 genes of a known defense gene), prophages, plasmids, or some combination thereof for all, known defense, or predicted novel defense genes. Solid circles indicate the type of element genes reside in. Error bars represent the 95% confidence interval after resampling the 69 genomes 200 times.

Of the 2,168 proteins identified by DefensePredictor but not DefenseFinder, 1,309 were homologous to a known defense protein, but were not encoded with the genes they were previously found to function with. An additional 252 proteins were homologous to known defense proteins, but were only identifiable using the sensitive alignment tool HHblits (*23*). Finally, DefensePredictor identified 607 proteins that had undetectable homology to known defense proteins or for which their top alignment with a known defense protein had less than 50% coverage.

Clustering the set of ∼2,500 predicted defense proteins at 80% coverage and 30% identity and selecting one protein per cluster, revealed 782 unique proteins (Fig. 2C). This set included 142 homologs of known defense proteins encoded in their native systems, 320 homologs encoded in new contexts, 85 remote homologs, and 235 novel predicted defense proteins. We focus on these 235 novel predicted defense proteins for the remainder of our analyses (table S3).

We first cataloged the predicted domains of these novel proteins using HHblits to query the Protein family (Pfam) database. Of the 235 unique proteins, 129 (55%) contain a domain previously implicated in defense (table S3). These protein clusters were not categorized as homologs of known defense proteins because they had less than 50% coverage with a known defense protein. The two most common domains were P-loop NTPases and PD-(D/E)XK nucleases, found in 13 and 6%, respectively, of all unique predicted novel defense proteins (Fig. 2D). Strikingly, 45% of the novel predicted defense proteins had no detectable domains in common with known defense proteins, suggesting that they function via novel mechanisms.

Next, we investigated the genomic distribution and genomic context of the predicted novel defense proteins. We found that 91% of the unique predicted novel defense proteins had homologs encoded by five or fewer genomes in our *E. coli* strain collection, compared with 73% of known defense proteins, and 55% of all proteins (Fig. 2E). The enrichment of predicted novel defense genes in the accessory genome suggests that they are often mobile and horizontally transferred, like known defense genes (*24*, *25*).

To examine the frequency with which the predicted novel defense genes reside in mobile genetic elements, we identified all prophages and plasmids in our *E. coli* collection using geNomad (*26*). Considering all proteins individually (*i.e.*, no longer grouping by protein clusters), we found that 20% of the predicted novel defense genes reside in plasmids, compared with 26% of known defense genes and 8% of all genes (Fig. 2F). Similarly, we found that 24% of predicted novel defense genes reside in predicted prophages, compared with 15% of known defense genes and 6% of all genes. Thus, the predicted novel defense genes reside in mobile elements well above background levels, similar to known defense genes. Finally, we found that 15% of predicted novel defense genes reside in defense islands (defined as within 10 genes of a known defense gene) compared with 17% of known defense genes and 2% of all genes (Fig. 2F). Thus, predicted novel defense genes are enriched in defense islands, though 85% are not within islands. Overall, the patterns of genomic distribution and context for our predicted novel defense genes resembles that of known defense genes.

### Predicted novel defense systems provide protection against phage

To assess whether the predicted novel defense genes in fact confer anti-phage defense, we selected those with a log-odds of defense > 0 (n = 2,264), and then extracted their putative transcriptional units (TUs) (Fig. 3A). We randomly selected 47 TUs for validation. We selected 59 additional TUs based on properties of interest, such as protein domains not previously validated in defense, novel domain combinations, or particularly high probabilities of defense, resulting in a total of 106 TUs selected for validation (table S4).

**Figure 3.**
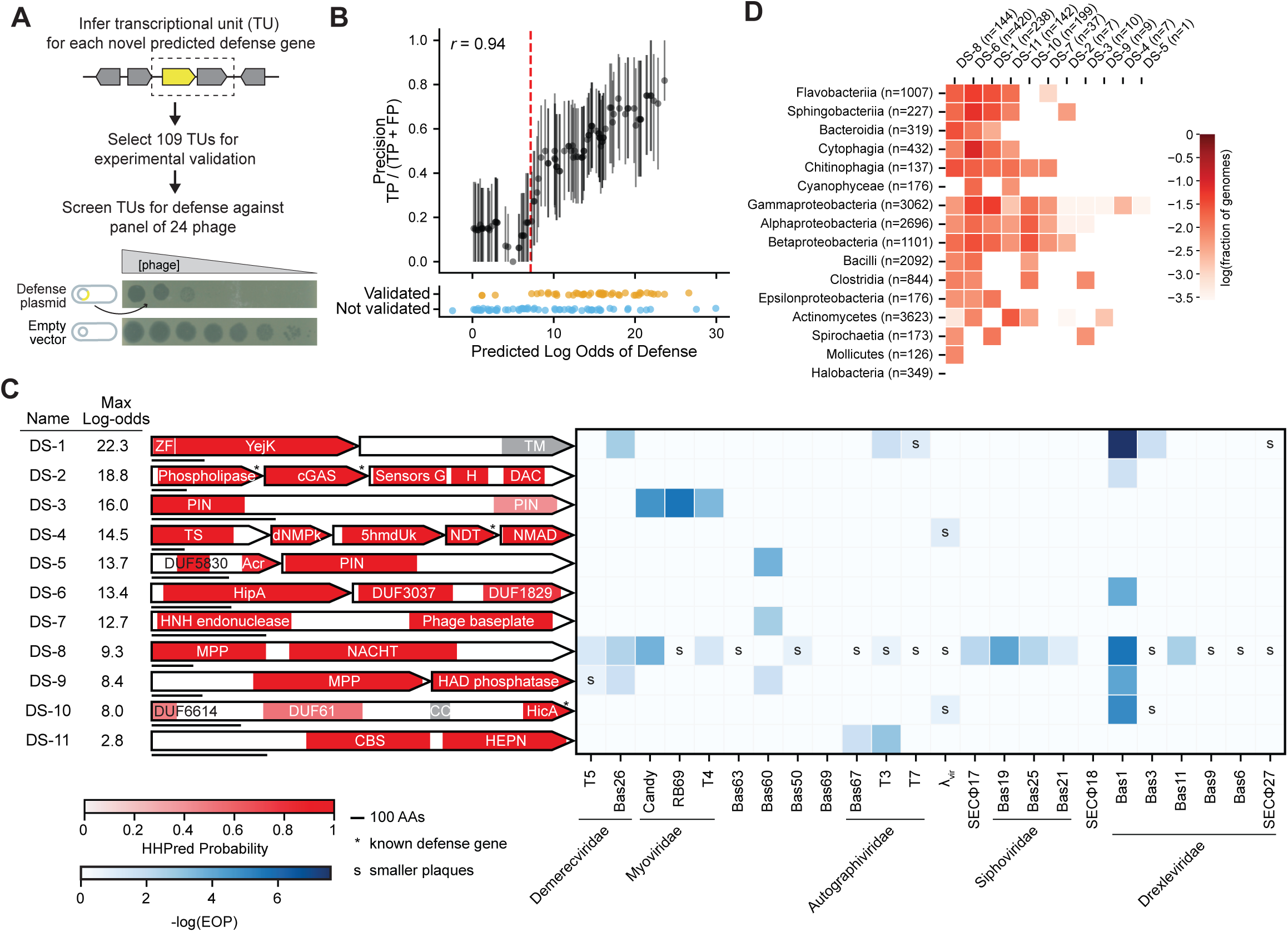
Predicted defense systems validate experimentally at a high rate. (**A**) Pipeline for validating predicted defense systems. The yellow gene represents a predicted novel defense gene. The boxed genes represent a putative transcriptional unit. An example plaquing assay is shown, with phage plaques forming on a lawn of *E. coli*, at decreasing concentrations of phage. (**B**) Bottom: each point represents one predicted system. Systems are plotted by their log-odds of defense, and colored by whether they provided protection. Top: validation rate at increasing log-odds thresholds, estimated by taking all systems `±`2 log-odds of each point. Error bars represent 95% confidence intervals after resampling systems 1,000 times. Pearson correlation between the log-odds and estimated validation rate is indicated. Red dashed line represents a log-odds cutoff of 7.2. TP, true positive; FP, false positive. (**C**) Left: domain annotations for validated defense systems containing domains with a defined function that had not previously been validated in defense. Horizontal bar underneath each system represents 100 amino acids. Genes with an asterisk represent homologs of known defense genes. The name and maximum log-odds of each system is indicated. Right: defense heatmap with EOP differences indicated by blue squares and plaque size differences indicated by the letter “s”. Phages are ordered by taxonomy. ZF, zinc finger; DAC, di-adenylate cyclase; TS, thymidylate synthase; dNMPk, dNMP kinase; 5hmdUk, 5-hmdU DNA kinase; NMAD, nucleotide-modification associated domain; MPP, metallophosphatase; TM, transmembrane; CC, coiled-coil; AA, amino acid. (**D**) Taxonomic distribution of systems with novel domains. The frequency of each system having homologs across the 16 most abundant prokaryotic classes is indicated by red squares.

To test for anti-phage defense, we placed each TU with its native promoter region on a low-copy number plasmid in *E. coli* MG1655 and then challenged these strains with a panel of 24 diverse *E. coli* phages (fig. S2). In total, 45 (42% of 106) of the cloned TUs produced smaller plaque sizes or reduced the efficiency of plating (EOP) at least ten-fold relative to an empty vector control strain (Fig. 3B, table S5). We refer to these validated TUs as DefensePredictor discovered Systems (DSs), with genes in multi-gene TUs denoted by an alphabetical suffix, e.g., *DS-8A* is the first gene of DS-8.

To estimate the experimental validation rate at different predicted thresholds of defense, we first defined the log-odds of each TU as the maximum log-odds of its constituent genes. We then calculated the validation rate for groups of TUs that were within four log-odds of each other. We observed a strong correlation between the validation rates and predicted log-odds of defense *(r =* 0.94, Fig. 3B), suggesting that DefensePredictor can effectively discriminate between defensive and non-defensive TUs.

To begin elucidating the function of the 45 validated TUs, we further annotated their protein domains (see Methods; table S6). This analysis identified five Pfam Domains of Unknown Function that were newly implicated in defense: DUF3037, DUF1829, DUF5830, DUF6988, and DUF6650. We also found 10 domains with a defined function that had not previously been validated in defense, including a YejK nucleoid-associated domain, a di-adenylate cyclase, a PIN nuclease, a 5-hydroxymethyluracil kinase, a thymidylate synthase, a HipA-family kinase, a phage baseplate protein, a metallophosphatase, a HicA mRNA interferase, and a CBS adenosyl-binding/dimerization domain (Fig. 3C; N.B., the PIN domain was recently validated elsewhere (*10*, *27*), but after DefensePredictor was run so it was not in the training set). We searched for homologs of the 11 systems that contain these novel defense domains in the ∼17,000 representative genomes and found that 9 had homologs in at least two bacterial classes (Fig. 3D), indicating that these systems, which likely function via novel mechanisms, are broadly distributed in bacteria.

For the 34 additional defense systems that we validated, 13 have PD-(D/E)XK nuclease domains, 7 have P-loop NTPase domains (e.g. AAA+, NACHT, ABC, etc.), and 4 have HEPN nuclease domains (Fig. 4A), though each was embedded within a protein or operon that distinguished it from known defense systems that also harbor these domains. Of the 34 systems containing either known defense or uncharacterized domains, 23 had homologs outside the proteobacteria, with three systems (DS-41, DS-26, and DS-29) even present in the archaeal Halobacteria (Fig. 4B). Thus, many of these systems have homologs that likely provide protection in a broad range of species.

**Figure 4.**
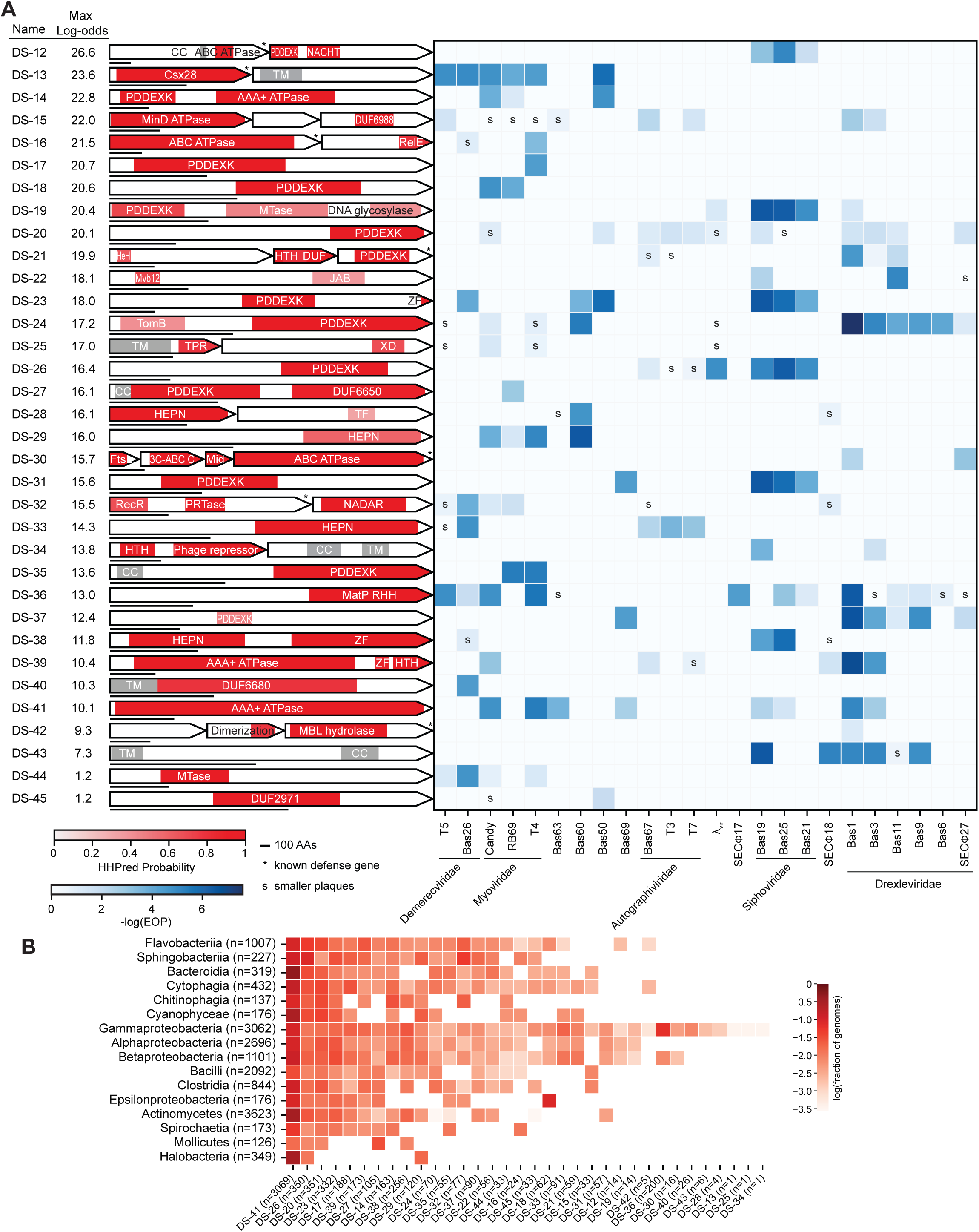
Validated systems contain diverse domains in diverse contexts. (**A**) Defense heatmap (as in Fig. 3C) for validated systems with known defense domains or domains without a known function. MTase, methyltransferase; HTH, helix-turn-helix; HeH, helix-extension-helix; TPR, tetratricopeptide repeat; TF, transcription factor; PRTase, phosphoribosyltransferase. (**B**) Taxonomic distribution of systems from panel (**A**), presented as in Fig. 3D.

### Novel defense domains are essential for protection

To gain insight into the functions of novel systems identified by DefensePredictor, we further investigated several systems containing domains not previously validated in defense. First, we investigated DS-8, a single protein system containing metallophosphatase and NACHT domains (Fig. 5A). Notably, the metallophosphatase domain is homologous to the human protein SMPDL3A, a phosphodiesterase that cleaves cGAMP to modulate cGAS-STING immunity signaling (*28*). The AlphaFold3-predicted structure of DS-8 (figs. S3A-B) closely aligned with the solved structure of SMPDL3A (*29*) (Fig. 5B). Homologs of the NACHT domain, a P-loop NTPase, feature in other defense systems (*6*, *30*, *31*) where they typically facilitate oligomerization and recognition of phage proteins, leading to activation of an N-terminal effector. Mutating either of two predicted catalytic residues in the metallophosphatase domain or a residue likely critical to NACHT ATPase activity completely ablated defense by DS-8, indicating that both domains and metallophosphatase activity are essential for protection (Fig. 5A, table S5). The identity of the cyclic nucleotide degraded by DS-8 remains to be identified, but this system highlights a new role for metallophosphatases in anti-phage defense and, given the homology to SMPDL3A, reveals a new connection between prokaryotic and human immunity.

**Figure 5.**
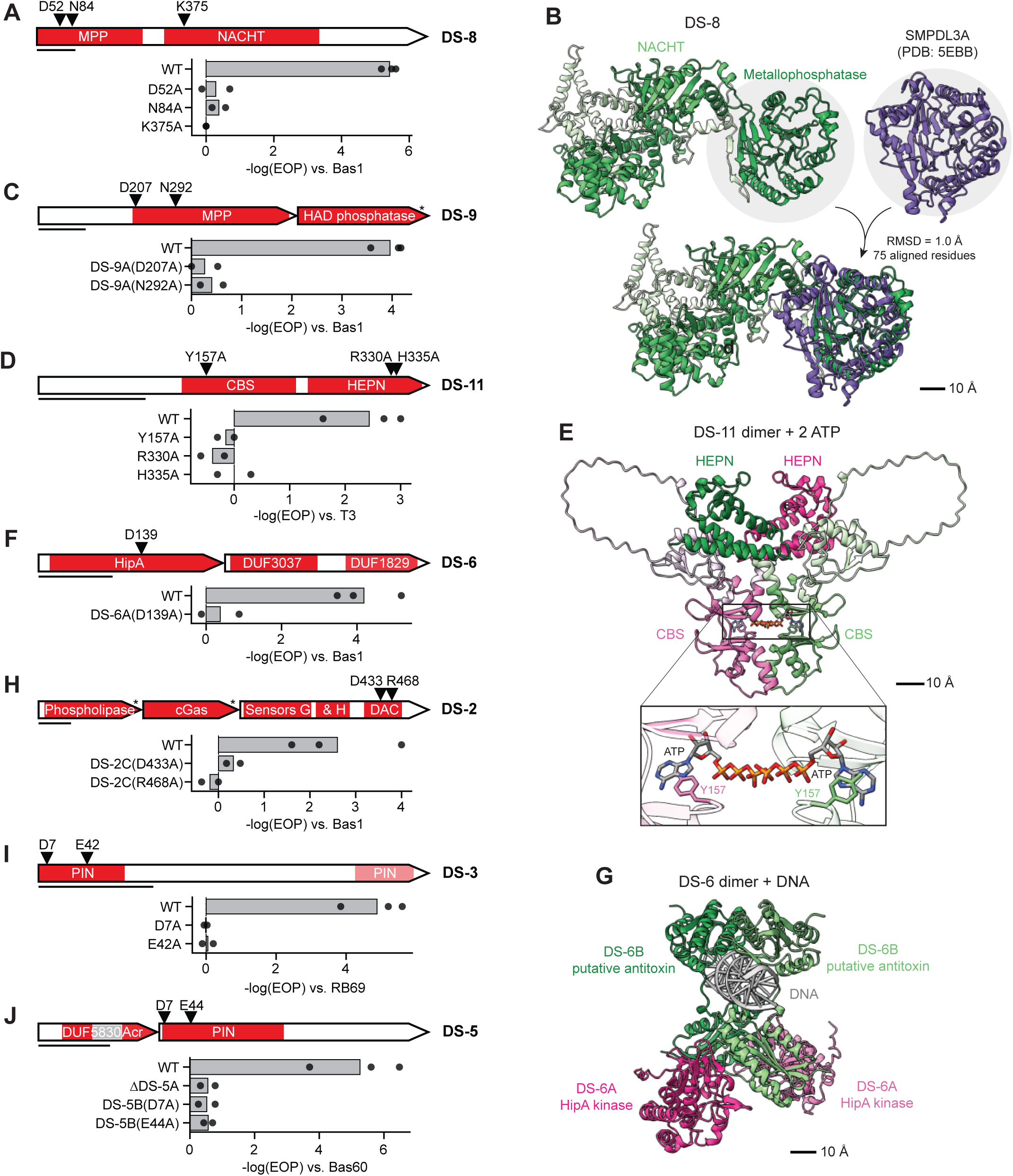
Novel defense domains are essential for protection. (**A**, **C**, **D**, **F**, **H**-**J**) For the systems DS-8 (**A**), DS-9 (**B**), DS-11 (**D**), DS-6 (**F**), DS-2 (**H**), DS-3 (**I**), DS-5 (**J**), the domain architecture is shown at the top along with the location of catalytic mutants tested. Bar graphs show -log(EOP) versus the phage indicated for the wild-type system and variants harboring the mutations indicated. Bars represent average values and points represent individual replicates. (**B**) Left: Predicted structure of DS-8 with domains highlighted in dark green. Right: solved structure of SMPDL3A (PDB: 5EBB). Bottom: Alignment between DS-8 and SMPDL3A. Scale bar is shown. (**E**) Top: predicted structure of a DS-11 dimer with two ATP molecules. One copy of DS-11 is shown in green and a second copy in pink. Protein domains are shown as dark regions. Bottom: predicted interaction between DS-11 protomers and ATP. The Y157 residues for are shown for each protomer. Scale bar is shown. (**G**) Predicted structure of two copies of DS-6A (pink), two copies of DS-6B (green), and DNA. Scale bar is shown.

The system DS-9 has two genes, the first harboring a metallophosphatase domain homologous to that of DS-8, and the second a predicted Haloacid Dehalogenase-like (HAD) phosphatase (Fig. 5C). HAD phosphatases are an enormous, but poorly understood family of enzymes that are often small molecule phosphatases (*32*). This domain has been identified in other defense systems (*9*, *33*), but its function in defense remains unknown. When we mutated the predicted catalytic residues in the metallophosphatase domain of DS-9A, we saw a loss of defense, suggesting it is essential for protection (Fig. 5C). As with DS-8, the target of the metallophosphatase remains unknown, as does that of the HAD phosphatase, but these two systems demonstrate that novel enzymes and novel signaling molecules likely remain to be discovered in anti-phage defense.

Next, we examined DS-11, a one-protein system containing CBS and HEPN domains (Fig. 5D). CBS domains facilitate dimerization of some enzymes by binding adenine derivatives including AMP, ATP, and Ap4A (*34*, *35*). There are no characterized defense systems with CBS domains, though these domains are prevalent in bacteria, archaea, and eukaryotes. HEPN domains are typically dimeric ribonucleases (*36*), including in known defense systems (*8*) such as the CRISPR-guided ribonuclease Cas13 (*37*). When we predicted the structure of two copies of DS-11 with two ATP molecules, AlphaFold3 predicted a reasonably strong interaction between DS-11 protomers (iPTM 0.59) and strong interactions between each protomer and ATP (iPTM 0.88) (Fig. 5E, figs. S3C-D). The predicted structure revealed a likely interaction between Y157 of the CBS domain of DS-11 and the adenine base of ATP. When we substituted Y157 with alanine, DS-11 no longer defended against phage (Fig. 5D), suggesting that binding of the CBS domain to an adenine nucleotide is essential to defense. Additionally, when we mutated catalytic residues in the HEPN domain, defense was ablated. Together, our data point to a model where the CBS domain of DS-11 regulates HEPN nuclease activity by binding an adenine derivative. Although many nucleotide signals have been implicated in phage defense, our validation of a CBS domain-based system suggests yet more remain to be found.

The two protein system, DS-6, features DS-6A, a predicted serine/threonine kinase related to the toxin HipA (*38*), and a second protein, DS-6B, with DUF3037 and DUF1829 (Fig. 5F) that has no similarity to HipA’s cognate antitoxin HipB. In MG1655, HipA phosphorylates glutamyl-tRNA synthase to block translation (*39*) unless neutralized by HipB, which also binds and represses the *hipAB* promoter (*40*). An anti-phage function for HipAB has not been reported. To determine whether DS-6B interacts with DS-6A and with DNA akin to HipB, we used AlphaFold3 to fold two copies of DS-6A with two copies of DS-6B, mimicking the stoichiometry of HipAB, and DNA (figs. S3E-F). AlphaFold3 predicted strong interactions between DS-6A and DS-6B (iPTM 0.76-0.79), and slightly weaker interactions between DS-6B and DNA (iPTM 0.67) (Fig. 5G). When we mutated a key predicted catalytic residue in DS-6A, we observed a loss of defense, suggesting that kinase activity is essential for protection. Thus, we hypothesize that DS-6 functions as a toxin-antitoxin (TA) system akin to HipAB, with the kinase activity of HipA providing protection against phage, possibly via an abortive infection mechanism similar to other TA-based defense systems (*41*, *42*).

The system DS-2 has three genes: a phospholipase, a cyclic GMP-AMP (cGAMP) synthase (cGAS), and a diadenylate cyclase (DAC) (Fig. 5H). The phospholipase and cGAS together constitute an apparent CBASS system (*43*) in which the cGAS presumably senses phage somehow, triggering cGAMP synthesis, which activates the phospholipase to degrade the cell membrane and abort infection. An association between CBASS and DACs was recently noted bioinformatically (*44*), but DACs have not been validated in phage defense. Using webFlaGs (*45*), we found that 15 of 30 homologs of the DAC identified here were co-located with a CBASS (fig. S4), confirming their association and suggesting they function together. To determine whether the DAC is essential for DS-2-based defense, we mutated two of its predicted catalytic residues. In each case, defense was eliminated. Thus, DS-2 may represent a novel type of CBASS system that employs two different cyclase genes.

DS-3 is a one-protein system with a PIN ribonuclease domain split between its N and C-terminus (Fig. 5I, figs. S5A-C). PIN domains are best characterized in VapC, the toxic component of VapBC TA systems, where they function to cleave tRNAs (*46*). When we mutated catalytic residues in DS-3, the system no longer defended against phage, suggesting this domain is essential for protection. There is no cognate antitoxin for DS-3, suggesting that it may be auto-inhibited until phage infection occurs.

Finally, DS-5 is a two protein system that also features a PIN ribonuclease, though not closely related to that of DS-3, along with a second protein containing a DUF5830 and a domain homologous to the anti-CRISPR protein, AcrIF4 (*47*). Mutating the catalytic residues in the PIN domain eliminated defense. Deleting the AcrIF4 domain-containing gene also ablated defense by DS-5 but did not lead to unrestrained toxicity of the ribonuclease indicating that DS-5 is likely not a TA system and that both components are essential to defense (Fig. 5J). All together, the systems probed here illustrate how DefensePredictor has identified novel anti-phage defense systems whose further study will likely reveal new mechanisms of immunity.

### Validated systems with known defense domains represent a reservoir of novelty

Although systems with domains not previously validated in defense represent the most obvious source of novelty, many of the systems with known defense domains also have the potential to function via novel mechanisms. For example, when we compared DS-38, which contains a HEPN and zinc-finger domain, to the known defense protein PD-T7-4, which contains these same domains, we saw strong alignment only of their HEPN domains (figs. S6A-E), suggesting that they use their Zn-finger domains in different ways and function via distinct mechanisms.

We also compared the predicted structures of all 14 PD-(D/E)XK nuclease proteins from our validated systems with the solved structure of the well-characterized restriction enzyme EcoRI (*48*). There was strong alignment near each protein’s PD-(D/E)XK catalytic motif (figs. S7A-B), but extensive structural diversity outside this catalytic core (figs. S7C & S8). For example, several proteins had large additional domains distal from the catalytic core (e.g. DS-12B, DS-14, DS-19, etc.). As with DS-38, the structural diversity of these proteins suggests they function via distinct mechanisms.

The system DS-32 has two genes: a NADAR domain-containing gene, and a phosphoribosyltransferase (PRTase). Despite low sequence similarity (16% identity across 63 aligned residues), the NADAR domain-containing protein DS-32B showed strong structural similarity to the antitoxin DarG, an ADP-ribosylglycohydrolase, from the TA system DarTG (*42*) (figs. S9A-E). However, the PRTase encoded by DS-32A and the ADP-ribosyltransferase DarT showed no similarity at the structural or sequence levels (figs. S9A and S9F-I). Thus, these systems likely function via distinct mechanisms despite the structural homology between their NADAR proteins, and may represent another example of TA partner swapping (*49*, *50*). Taken together, these examples illustrate how even systems identified here with known defense domains have the potential to function via novel mechanisms.

### Thousands of predicted novel defense genes remain to be validated in *E. coli* and beyond

We sought to assess the landscape of defense systems in the broad *E. coli* pangenome, beyond the initial set of 69 strains examined. We therefore applied DefensePredictor to a large, diverse set of 3,000 *E. coli* and *Shigella* genomes. We clustered the proteins encoded by these genomes at 80% coverage and 30% identity, and saw that most genes reside in the accessory pangenome, with 80% present in 10 strains or fewer (Fig. 6A). Even with 3,000 strains, the number of unique protein clusters was not saturated, underscoring the enormous diversity of the *E. coli* pangenome (fig. S10A). We ran DefensePredictor on all 3,000 strains and found an average of 28 defense systems per genome, compared with an average of 4.5 for DefenseFinder (Fig. 6B). We found that DefensePredictor output similar predictions for homologs, where 82% of clusters with at least one predicted defense protein had a majority of proteins in the cluster predicted as defensive (fig. S10B). Thus, we plotted the number of unique protein clusters containing at least one predicted defense protein against the number of genomes included in our analysis (Fig. 6C). For each category of predicted defense (Fig. 2B), there was a non-saturating increase, with a total of 1,374 unique protein clusters harboring a predicted novel defense protein (table S8). The latest versions of DefenseFinder and PADLOC (*51*), another homology-based identification tool, identified only 56 (4%) and 48 (3%), respectively, of these unique proteins. Together, our results indicate that we still have only just scratched the surface of defense system diversity in the *E. coli* pangenome.

**Figure 6.**
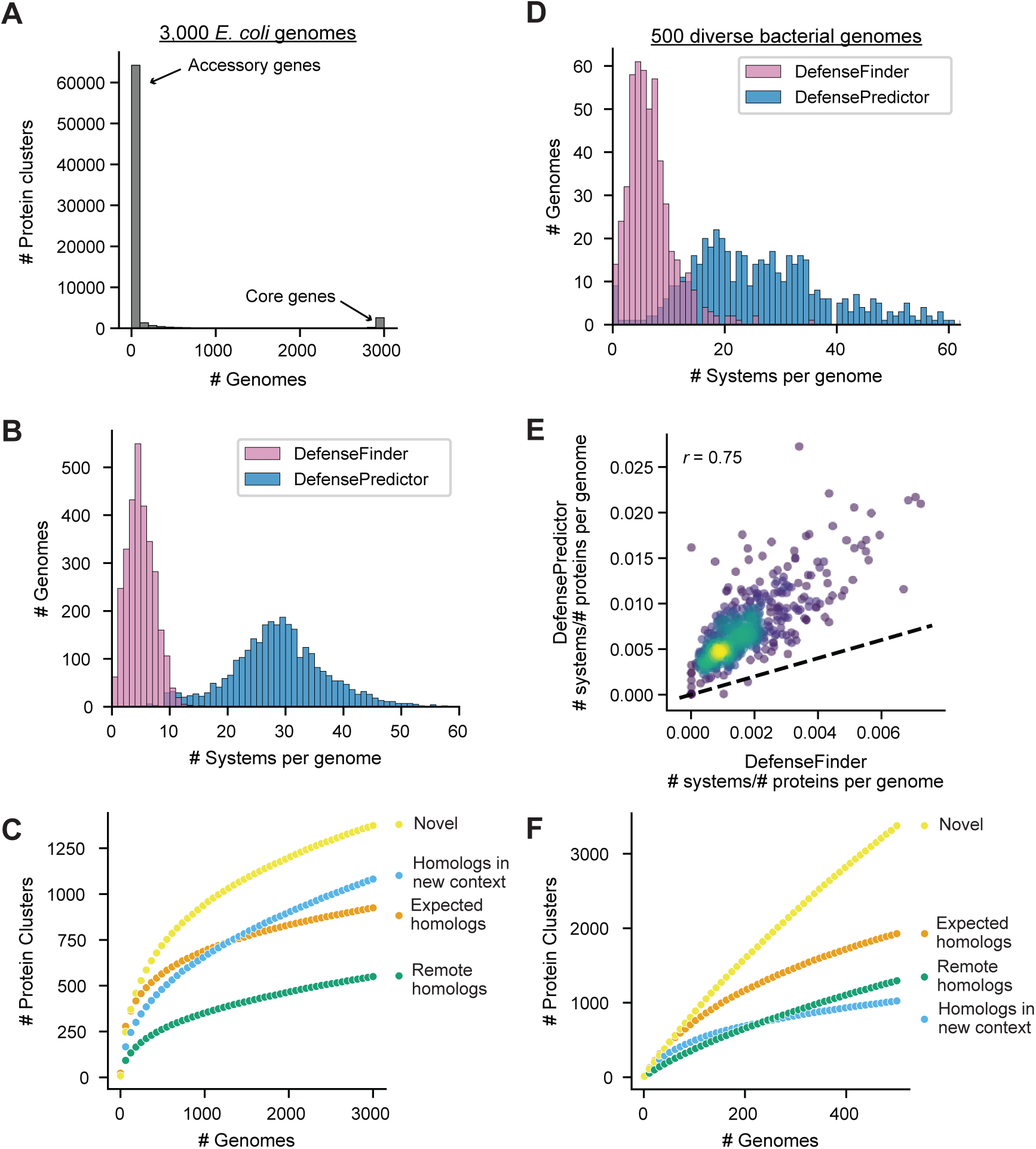
Thousands of predicted defense proteins remain to be validated. (**A**) Histogram of the number of genomes each protein cluster is encoded by in a set of 3,000 *E. coli* strains. (**B**) Distribution of the number of systems identified by DefenseFinder or DefensePredictor in a set of 3,000 *E. coli* strains. (**C**) Average number of unique protein clusters for different categories of predicted defense proteins (see Fig. 2B) when considering between 1 and 3,000 *E. coli* genomes in steps of 50. Averages were calculated after resampling genomes 50 times at each step. (**D**) Same as (**B**) for 500 diverse bacterial strains. (**E**) Comparison of the number of systems identified by DefenseFinder or DefensePredictor, normalized by the number of genes for each genome. The dashed line represents the function *y* = *x*. Pearson correlation is indicated. (**F**) Same as (**C**) for 500 diverse bacterial strains.

Finally, to explore the defense landscape outside of *E. coli*, we analyzed 500 randomly selected genomes from the set of ∼17,000 representatives. The ∼300,000 unique protein clusters encoded by these 500 genomes (figs. S10C-D) far exceeded the ∼70,000 unique protein clusters encoded by the 3,000 *E. coli* strains. We ran DefensePredictor on all 500 genomes and found an average of 27 defense systems per genome, compared with an average of 6.2 for DefenseFinder (Fig. 6D). Both approaches agreed on which genomes were enriched with defense systems (Fig. 6E), hinting that DefensePredictor may identify more systems in bacteria that experience greater phage predation, as evidenced by the enrichment for known systems in those bacteria. When we plotted the number of unique predicted defense proteins against the number of genomes included in our analysis, we again saw a non-saturating increase, with a total of 3,376 unique protein clusters harboring a predicted novel defense protein (table S9). These results suggest that DefensePredictor will enable the discovery of scores of additional systems in diverse bacteria.

## Discussion

DefensePredictor takes as input the sequence of a gene and its genomic neighbors, and predicts whether this gene functions in anti-phage defense. When applied to a set of 69 diverse *E. coli* strains, we predicted over 700 unique defense proteins, including >200 with no detectable homology to or less than 50% coverage with any known defense protein. For this latter set of >200, we achieved an experimental validation rate of 42% with a strong correlation between the predicted log-odds of defense and the experimental validation rate. The new systems we identified indicate that *E. coli* harbors a much broader landscape of anti-phage defense than previously appreciated, expanding the likely number of systems by multiple orders of magnitude. The new systems feature domains not previously validated in defense, including homologs of eukaryotic innate immune proteins.

Although some predicted defense systems did not validate and could represent false positives, we anticipate that many will validate if challenged with a larger, more diverse panel of phages. Further, some systems may not be adequately expressed in *E. coli* K-12 strains and may function in a different host strain. Finally, detecting the defense function of some systems may require redefining the boundaries of the system to capture additional genes or regulatory elements.

The *in-silico* test results suggest that DefensePredictor has a false negative rate of ∼25%. This can likely be improved by including the dozens of systems discovered here and elsewhere since the start this work into a new training set. New PLMs (e.g. see refs (*52*, *53*)) may also improve model performance. DefensePredictor did not identify all of the genes identified strictly through homology searches, *e.g.* via DefenseFinder, suggesting that both approaches, and experimental screening, will be needed to comprehensively catalogue the repertoire of defense systems in a strain of interest. Finally, we note that the training set used for DefensePredictor features a predominance of systems identified or validated in *E. coli*, so whether the model performs as well on other species, particularly those distantly related to *E. coli*, remains to be tested.

Application of DefensePredictor to a set of 3,000 *E. coli* and 500 diverse bacterial genomes indicated that the number of predicted systems does not saturate. This finding underscores both the enormous genetic diversity of even a single species of bacteria like *E. coli* and the vast number of anti-phage defense systems that remain to be found. As an open-source software tool that can be run on any prokaryotic genome of interest, we anticipate that DefensePredictor will help catalyze the mapping of such defense systems and the discovery of novel molecular functions, while also helping to further elucidate connections between prokaryotic and eukaryotic immunity.

## Methods

### Compute resources

Intel Xeon Platinum 8260 or Xeon Gold 6248 computers with Nvidia Volta V100 GPUs from MIT’s Supercloud (*54*) were used for computational analyses unless otherwise noted.

### Dataset assembly

17,454 assembled genomes were downloaded from RefSeq (*55*) on May 2, 2023, comprising all bacterial, archaeal, and viral representative and reference genomes. Genomes were searched for defense systems using DefenseFinder version 1.2.2, and all genes that were identified as part of a full system were labeled as our positive gene set.

To construct the negative gene set, GO Processes were extracted from the General Feature Format (GFF) table for each genome. GO processes were deemed non-defensive if more than 60% of gene members were not identified by DefenseFinder. Only six GO processes did not meet this threshold: ‘maintenance of CRISPR repeat elements’, ‘DNA modification’, ‘DNA methylation’, ‘DNA restriction-modification system’, ‘defense response to virus’, and ‘nucleic acid phosphodiester bond hydrolysis’. Genes belonging to COGs of mobile elements as defined by Marakova et al. (*4*) were added to the negative set as well. Finally, homologs of secretion system effectors from BastionHub (*56*) were identified with MMseqs2 with coverage and identity cutoffs of 90% and added to the negative gene set. Genes were removed from the negative set if they were identified by DefenseFinder or if they were part of a TA system as indicated by their name containing one of the phrases ‘toxin-antitoxin’, ‘addiction module’, or ‘abortive infection.’

### Model training

To generate ESM2 embeddings for the feature matrix the 150 million parameter ESM2 model (*57*), ‘esm2_t30_150M_UR50D’, was used. The feature matrix was then assembled as described in the Results. The LightGBM (*58*) framework was used to train a gradient boosting classifier with the feature matrix. Hyperparameters for the classifier were fit with optuna (*59*), using the validation data as a held-out set. The number of leaves for the model was fit between 4 and 256, and the minimum number of child samples was fit between 32 and 1,024 for 20 trials. The final model used the selected parameters of 255 leaves and 782 minimum child samples. We also set the learning rate to 0.01 and number of estimators to 100,000 for training. Training was stopped after the average precision on the validation data did not increase for 200 iterations.

### Calculation of precision and recall

To calculate the precision and recall of each modeling approach, test genes were ranked by probability, p-value, e-value, or cosine similarity for DefensePredictor, guilt-by-association, HMM search, or ESM cosine similarity, respectively. To prevent these metrics from being dominated by the largest clusters of defense genes or the largest GO process or COGs, precision and recall for each test gene was weighted such that the sum of weights for each of these groups equaled one. More formally, at each threshold, *p*, we calculated precision as 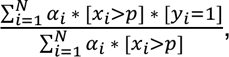 and recall as 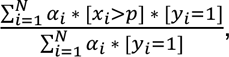 such that 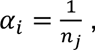 where *N* is the number of genes in the test set, *α_i_* is the weight for sample *i*, *x*_*i*_ is the predicted confidence of defense for sample *i*, *y_i_* is the label for sample *i*, and *n* is the size each gene group *j*, such that *i* ∈ *j*.

### Guilt-by-association implementation

To implement a guilt-by-association approach, all proteins in the set of representative genomes that belong to an MMseqs2 cluster containing at least one protein in the test set (positively or negatively labeled) were extracted. For each gene that codes for one of these proteins, whether it resides within 10 genes of a DefenseFinder-identified defense gene was assessed. In making this assessment, neighboring defense genes belonging to an HMM cluster in the test set or that were part of a known system with the query gene were discarded. Then for each MMseqs2 cluster, the significance of its association with known defense genes was calculated using Fisher’s exact test, where the frequency that each MMseqs2 cluster resided within 10 genes of a defense gene was compared with random expectation. All test genes were ranked based on the significance of their cluster’s association.

### SHAP value calculation

The contribution of each feature to the final output prediction of each gene in the test set was calculated using the shap package (*60*). To identify the most important individual features, SHAP values were aggregated for each feature across all true positive genes in the test set using a weighted average, such that each HMM cluster contributed equally to each feature’s average.

The contribution of each neighboring gene was calculated by summing the SHAP values for all of its features. After summing SHAP values for each gene neighbor, the gene neighbors that were defensive were identified using DefenseFinder, and the gene neighbors that were transposases or integrases were identified by searching for genes with ‘transposase’ or ‘integrase’ in their name, respectively.

### HHblits searches

To search with HHblits, HMMs for query proteins were built by searching the UniRef30_2019_11 database with two iterations of HHblits. These built HMMs were then used to search the Pfam35 (*61*) database from hh-suite (*62*), or a custom-built database of known defense proteins. To build a database of HHblits-compatible HMMs of known defense proteins, the UniRef50 (*63*) dataset was searched for homologs of each known defense protein using HMMER (*64*). The sequence with the highest score was selected for each known defense protein, and these sequences were used to search the UniRef30_2019_11 database with two iterations of HHblits to build HMMs.

### Selection of defense systems for experimental validation

TUs for each predicted defense gene were defined by taking groups of consecutive genes, transcribed in the same direction, and each separated by 30 base pairs (bps) or less. To capture the native promoter of each TU, we included 200 bp upstream of the 5’-most gene, or until the next gene was reached if it was closer. To capture the terminator of each TU, we included 100 bp downstream of the 3’-most gene, or, again, until the next gene if it was closer.

47 TUs were randomly selected for validation, such that each TU had at least one predicted novel defense gene that was not homologous to the other selected genes. 28 TUs were selected for having a high log-odds of defense. 17 TUs were selected for the presence of novel domains. Finally, 14 TUs were hand selected based on domain annotations.

Six selected TUs were not cloned because they were found in type VI secretion system regions and had homology to known effector toxin or immunity proteins (table S4), likely representing false positives. In our calculation of validation rate these TUs were counted as unvalidated.

### Phage taxonomy construction

The phage phylogenetic tree was built using VICTOR (*65*). Phage genomes from NCBI were compared at the DNA level. Phage taxonomy was built using VICTOR’s D0 algorithm, which compares phages based on the length of all homologous regions divided by genome length. Phages for screening were selected to maximize phylogenetic diversity.

### Bacterial and phage culture conditions

*E. coli* cells were grown in Luria Broth (LB) at 37 °C unless otherwise noted. Select ECOR strains were obtained from the Thomas S. Whittam STEC Center at Michigan State University (*22*), and UMB isolates were obtained from Alan J. Wolfe at Loyola University Chicago (*66*).

Phages were propagated overnight from single plaques. To harvest phages, cultures were treated with chloroform, spun down, and the resulting supernatant was extracted. The identity of each phage was confirmed by PCR (primers in table S10). Phages were stored at 4 °C.

Phage phiLS50-11 (referred to here as Candy) was isolated during Harvard’s LS50 course in the fall of 2021 by students under the instruction of S. Srikant. It was isolated from Charles River water on *E. coli* MG1655, and sequenced to taxonomically classify it as *Tevenvirinae*.

### Strain construction

TUs were PCR amplified from their native strains (table S10). Gibson assembly was used to insert defense systems into pCV1 (*8*), which has a pSC101 origin of replication, chloramphenicol resistance cassette, and no promoters upstream of the insertion site. One system, DS-44, was cloned into pBAD, which has an upstream arabinose-inducible promoter. Primers for DS-44 were designed to avoid an upstream integrase and thus an inducible promoter was necessary to drive expression. Plasmids were verified by whole plasmid sequencing and transformed into *E. coli* MG1655 for screening.

Catalytic residues were identified by aligning with previously characterized proteins. Mutations to catalytic residues were generated using site-directed mutagenesis. For each catalytic mutant, completely overlapping PCR primers facing opposite directions and containing the desired mutation were designed. Each PCR was run for 18 cycles, and the linearized product was transformed into *E. coli* DH5α cells. Re-circularized products were verified by whole plasmid sequencing and transformed into *E. coli* MG1655. The same process was used to delete plasmid regions, except outward-facing primers were designed flanking each region-to-be-deleted and each primer contained homology at their 5’ end to the other side of the region.

### Efficiency of plating assays

To test for anti-phage defense, strains were grown overnight in LB, diluted 1:100 into melted LB + 0.5% agar, plated onto LB + 2% agar, and left for 30 minutes at 22 °C. Eight 10-fold serial dilutions were generated for each phage, and 2 μL of each dilution was dispensed onto *E. coli* lawns. Plates were incubated overnight at 37 °C. Plaque forming units (PFUs) were measured by counting the number of individual plaques at the most concentrated dilution where such plaques were discernable and multiplying this count by the dilution factor (table S5). In instances where individual plaques bled into each other, the outer contours of plaques were used to count. If plaques could not be discerned at the lowest visible dilution, then a default count of 50 was used. EOP for each predicted defense system was calculated by dividing its PFU by the PFU of an empty vector strain.

TUs that did not validate in LB at 37 °C from the set of 48 randomly selected TUs were screened in slow growth conditions, where LB medium was replaced with M9L medium (M9 salts supplemented with 0.2 mM CaCl2, 2 mM MgSO4, 0.1% casamino acids, and 0.4% glycerol), and cells were grown at 30 °C. The slow growth condition yielded two new positive hits (DS-30 & DS-36).

### Annotating validated defense systems

Domains for validated proteins were comprehensively annotated using the web tool HHpred (*67*) to search the Pfam database, the Protein Data Bank (PDB) (*68*), the Structural Classification of Proteins (SCOPe) database (*69*), and the Conserved Domain Database (CDD) (*70*). HHpred uses HHblits to carry out these searches, but it also incorporates information about each sequence’s predicted secondary structure, making it slightly more sensitive than the local HHblits searches. The highest probability domain was selected for each non-overlapping protein region. The label for each domain was curated based on all hits to the same region. Transmembrane regions of validated proteins were identified using DeepTMHMM (*71*). Coiled-coil regions were identified using DeepCoil2 (*72*). The PD- (D/E)XK motif was identified for each validated protein using HHPred alignments to characterized nucleases. The extracted PD-(D/E)XK motifs were aligned using Kalign (*73*) and visualized using JalView (*74*).

### Analyzing taxonomic distribution of validated systems

HMMs of each protein in a validated system were built by searching the UniRef30_2019_11 database with HHblits. HMMs were used to search for homologs across the ∼17,000 representative genomes using HMMER. Homologs were identified using an E-value cutoff of 0.001 and a coverage cutoff of 80% for query and target sequences. For multi-gene systems, system completeness was assessed by examining whether homologs of the same system were encoded next to each other in the genomes they were found in.

### Protein structure prediction and visualization

Protein structures were predicted using the AlphaFold3 server (*75*). Protein structures were visualized using ChimeraX (*76*). Structural alignments were generated using the matchmaker command in ChimeraX with default parameters. The predicted structures of the PD-(D/E)XK nucleases were aligned using the first aspartate in their catalytic motif. Most validated nucleases did not share enough homology to align using ChimeraX’s matchmaker.

### Predicting defense systems in set of 3,000 *E. coli* and 500 diverse bacterial strains

3,000 assembled *E. coli* or *Shigella* genomes were randomly selected and downloaded from NCBI. DefensePreidctor was run on all genomes. To calculate the number of systems identified by DefensePredictor, systems were defined as groups of consecutive genes, transcribed in the same direction, each separated by 30 bp or less, having at least one gene with a log-odds of defense greater than 7.2. Proteins were clustered at 80% coverage and 30% identity using MMseqs2. Protein clusters were categorized as in Fig. 2B, such that if any cluster member was an “expected homolog” this designation was given to the cluster, then “homolog in a new context,” then “remote homolog,” and finally “novel” if all predicted defensive cluster members were designated as such. The unique predicted novel defense proteins were searched for homologs using HMMER, with profiles from DefenseFinder 1.3.0 and PADLOC 2.0.0, using e-value and coverage cutoffs specified by each tool. Phage Defence Candidate (PDC) systems (*27*) were excluded from the PADLOC search because they are unvalidated. The 500 diverse bacterial strains were randomly selected from the set of ∼17,000 representatives and analyzed in the same manner.

## Supplementary Data

**Table S1.** Dataset for model development.

**Table S2.** SHAP values.

**Table S3.** Predictions for 69 diverse *E. coli* strains.

**Table S4.** TUs selected for screening.

**Table S5.** Plaquing assay quantification.

**Table S6.** Protein domains in validated systems.

**Table S7.** Summary statistics from AlphaFold3 predictions.

**Table S8.** Predictions for 3,000 *E. coli* strains.

**Table S9.** Predictions for 500 diverse bacterial genomes.

**Table S10.** Primer sequences and created strains.

## Code Availability

DefensePredictor is available at https://github.com/PeterDeWeirdt/defense_predictor

Code for all analyses is available at https://github.com/PeterDeWeirdt/defense_predictor_manuscript

## Acknowledgements

We thank A. Millman, C. Vassallo and S. Srikant for comments on the manuscript, and A. Keating, S. Ovchinnikov and C. Coley for feedback on the model development. This work was supported by a National Science Foundation Graduate Research fellowship to P.C.D. M.T.L. is an Investigator of the Howard Hughes Medical Institute.

## Author Contributions

M.T.L and P.C.D. designed the experiments and wrote the manuscript. P.C.D. designed and wrote code to train the model. P.C.D. analyzed data. P.C.D. and E.M.M. performed experiments and cloning.

---

**Figure S1.**
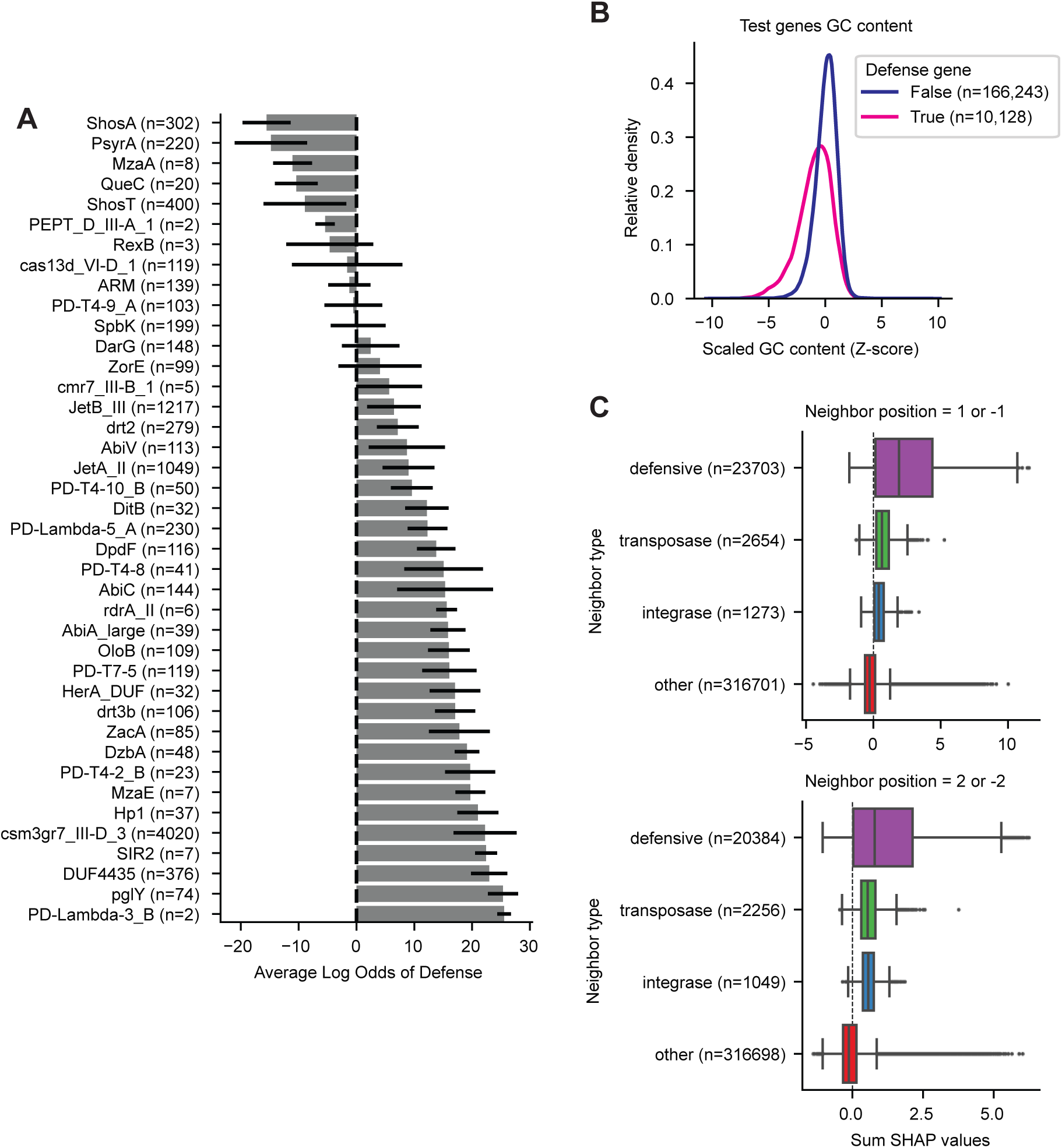
DefensePredictor leverages known genomic associations to make its predictions. (**A**) Average DefensePredictor log-odds for test genes grouped by HMM cluster. For clusters representing multiple HMMs, the HMM with the most connections to other HMMs was chosen as the label. Number of homologs for each cluster is indicated next to the cluster name. Error bars represent the standard deviations of predictions across homologs. (**B**) Z-scored GC content of genes in the test set, split based on which genes are defensive. Densities are not scaled between groups. The number of genes in each group is indicated. (**C**) Box plot of summed SHAP values for neighboring genes one or two away that are labeled as defensive, transposes, or integrases. The number of genes in each category is indicated.

**Figure S2.**
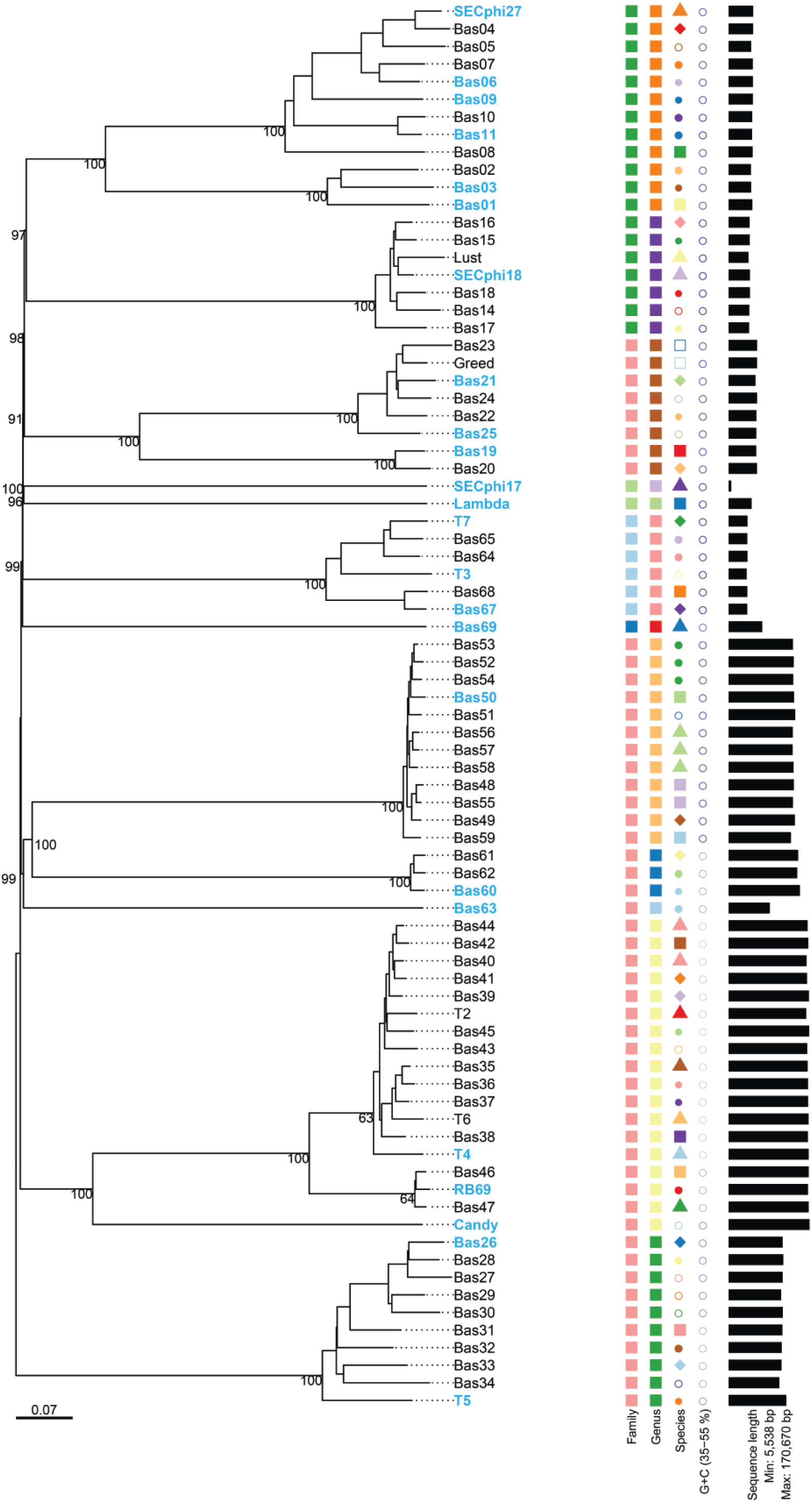
Selected phages represent a broad range of taxonomies. Phylogenetic tree of BASEL (*77*) and other common *E. coli* phage. Phages selected for screening are shown in blue. Family, genus, and species groups inferred from the tree are indicated. GC content is indicated, where low and high GC content genomes are represented by light gray and dark blue circles, respectively. Genome length is also indicated.

**Figure S3.**
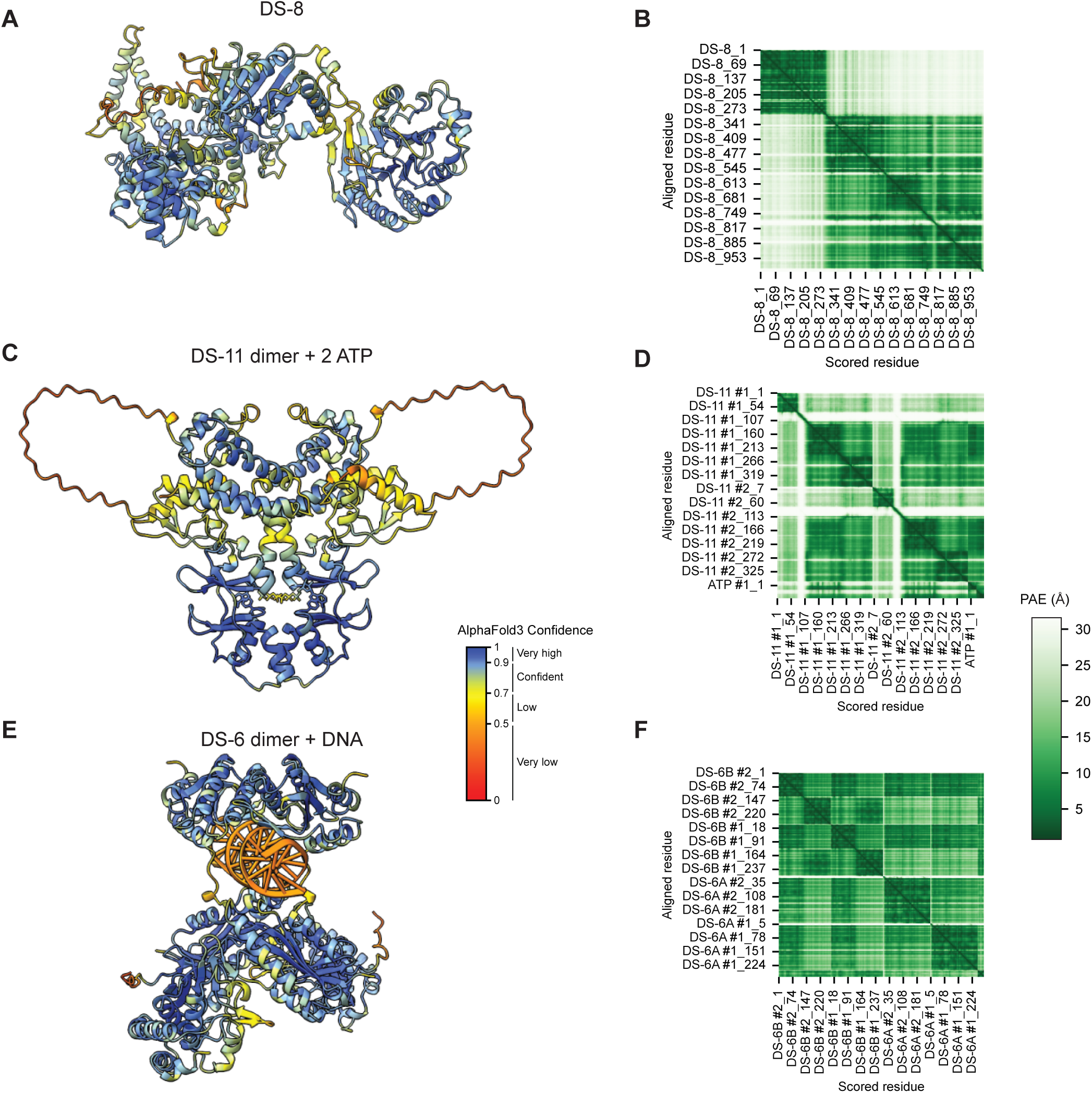
AlphaFold3 confidence metrics for select systems with novel defense domains. (**A**, **C**, **E**) Per-residue pLDDT confidence scores mapped onto the predicted structures for DS-8 (**A**), two copies each of DS-11 and ATP (**C**), two copies of DS-6 and DNA (**E**). (**B**, **D**, **F**) Predicted aligned error (PAE) between residues for DS-8 (**B**), two copies each of DS-11 and ATP (**D**), two copies of DS-6 and DNA (**F**).

**Figure S4.**
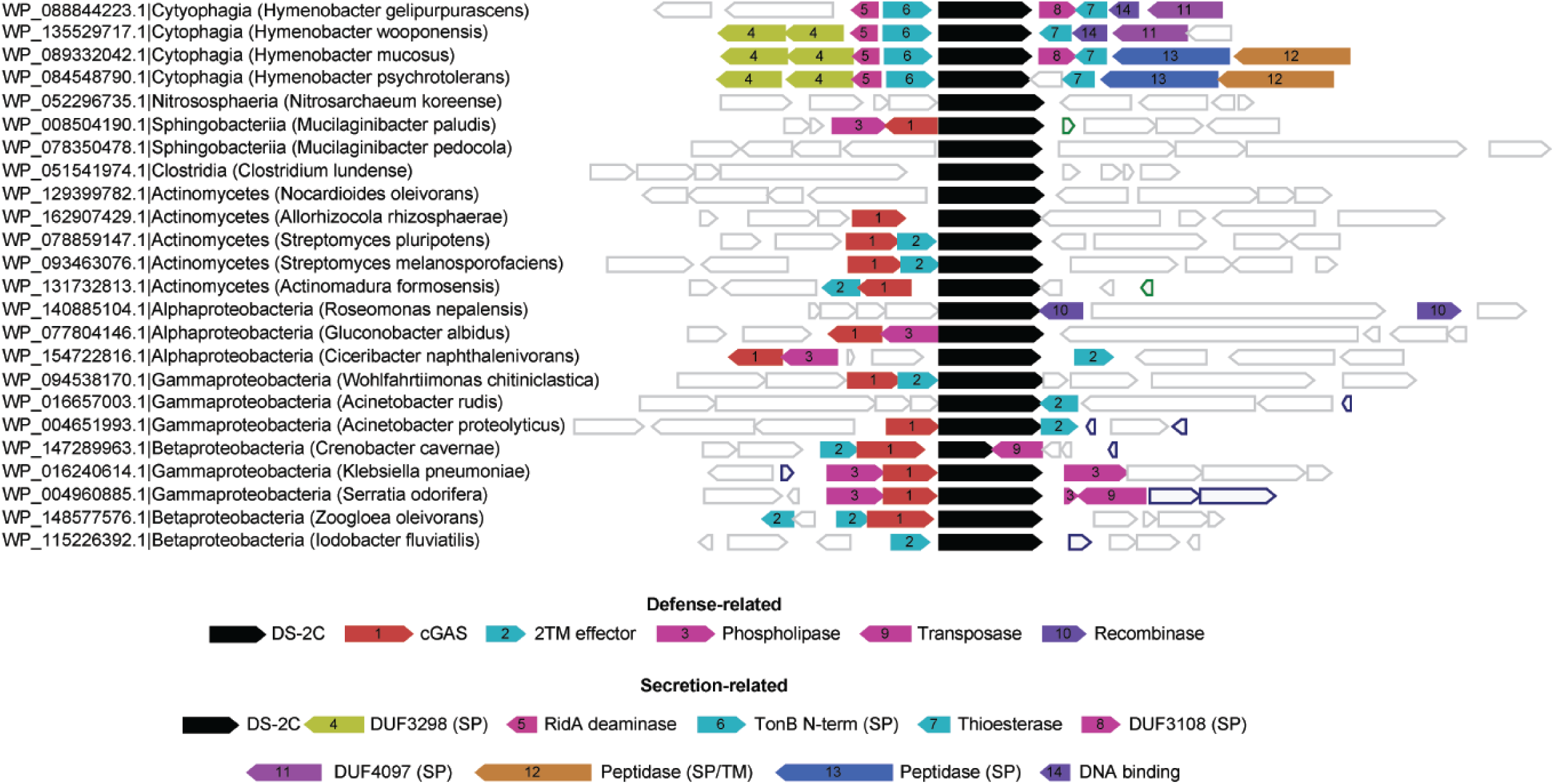
A di-adenylate cyclase co-occurs with CBASS genes. WebFlaGs output showing homologs of DS-2C and their genomic neighbors. The protein accession and species for each homolog is indicated. Clusters of neighboring genes share the same number are labeled at the bottom of the plot. SP, signal peptide-containing protein; TM, transmembrane domain-containing protein.

**Figure S5.**
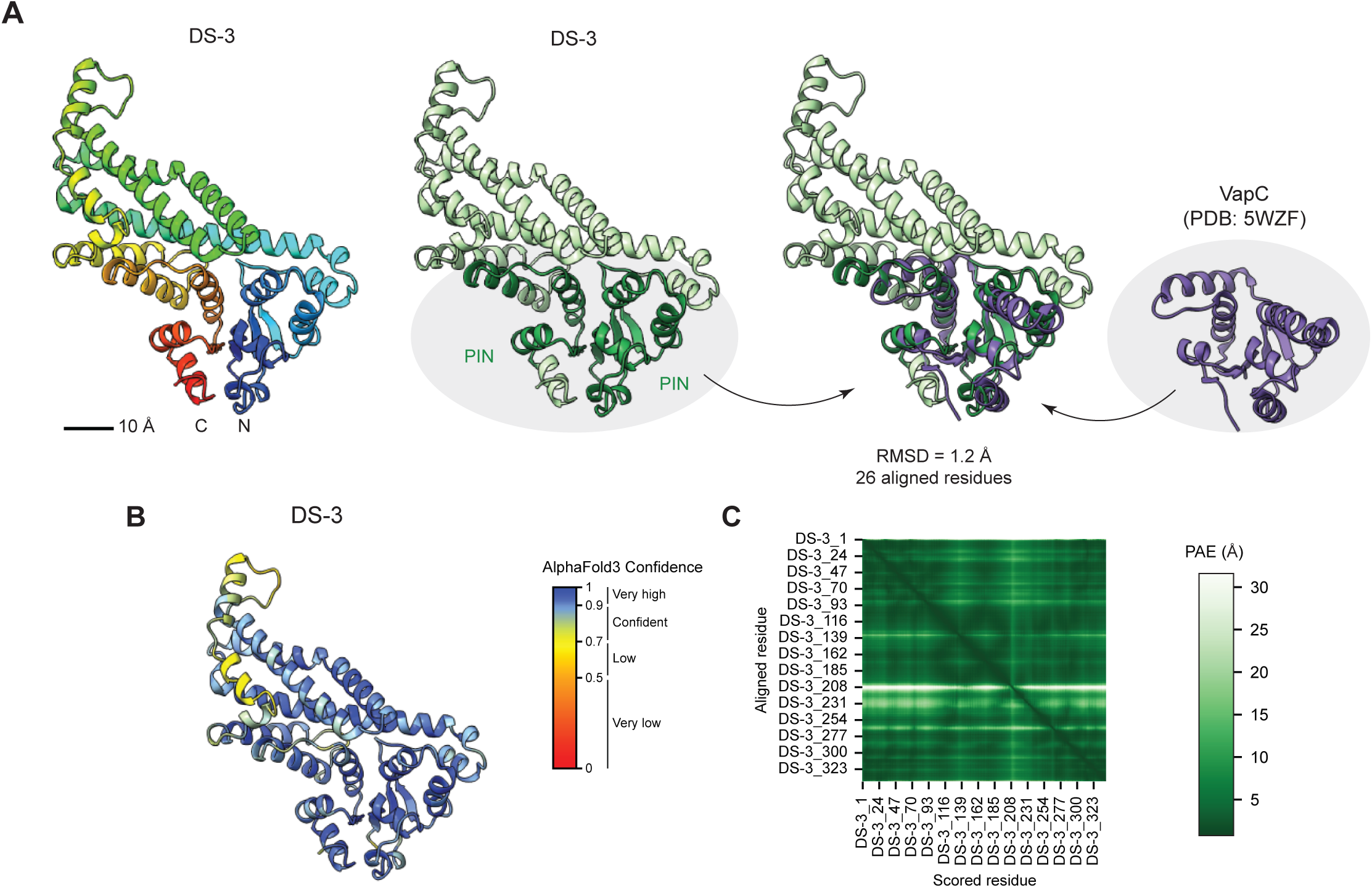
DS-3 has a PIN domain split between its N and C-terminus. (**A**) Predicted structure of DS-3, colored blue to red to indicate its N versus C-terminus, respectively (left), or with both halves of its PIN domain colored green (center). The predicted structure of DS-3 is compared with the solved structure of VapC from Mycobacterium tuberculosis (*78*). Scale bar is shown. (**B**) Per-residue pLDDT confidence scores mapped onto the predicted structure of DS-3 (**C**) Predicted aligned error (PAE) between DS-3 residues..

**Figure S6.**
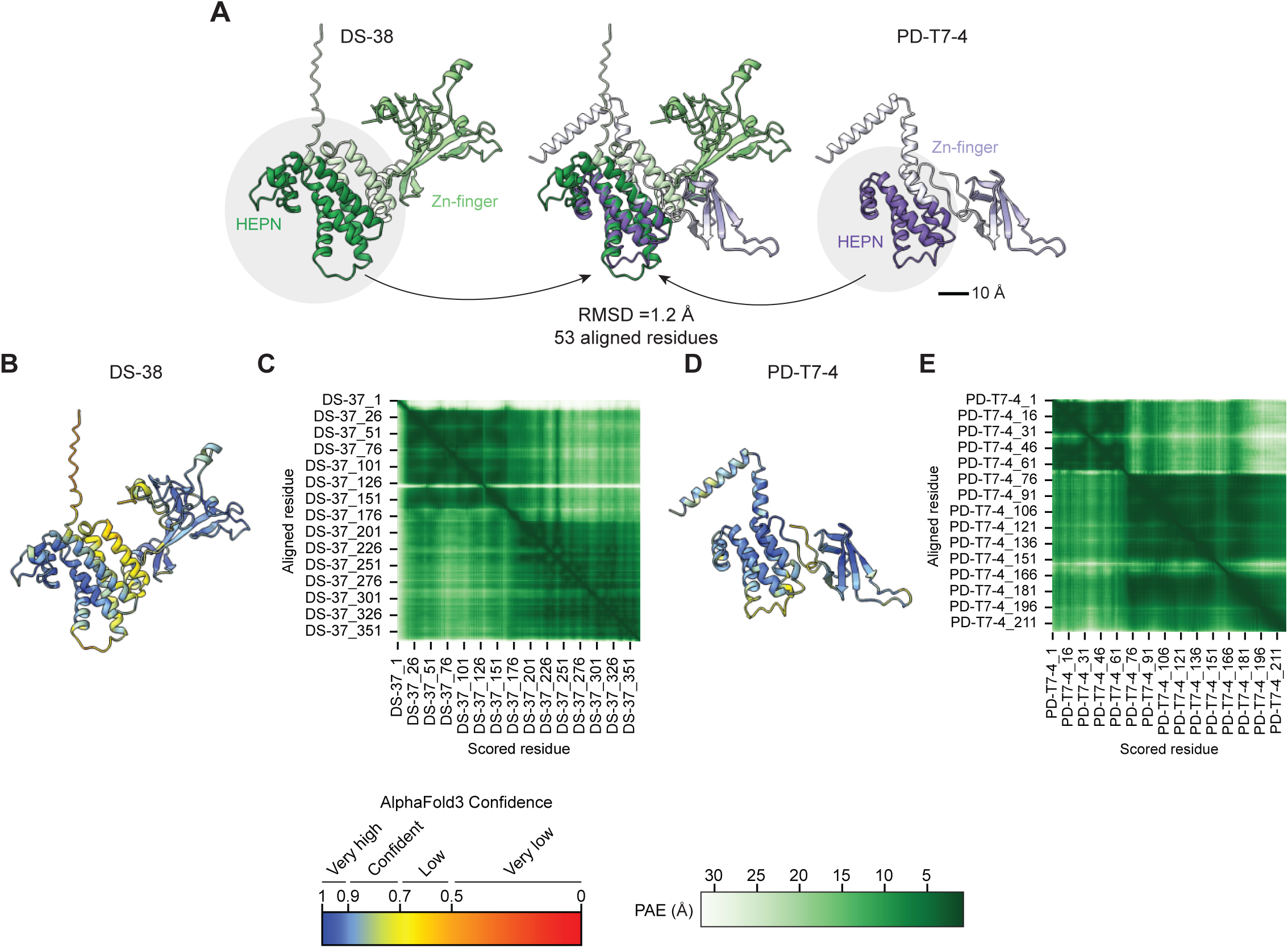
DS-38 and PD-T7-4 have distinct Zn-finger domains. (**A**) Left: predicted structure of DS-38 with domains annotated in dark green. Right: predicted structure of PD-T7-4 with domains annotated in dark blue. Center: Alignment between DS-38 and PD-T7-4. Scale bar is shown. (**B**, **D**) Per-residue pLDDT confidence scores mapped onto the predicted structures for DS-38 (**B**) and PD-T7-4 (**C**). (**C**, **E**) Predicted aligned error (PAE) between residues for DS-38 (**C**) and PD-T7-4 (**D**).

**Figure S7.**
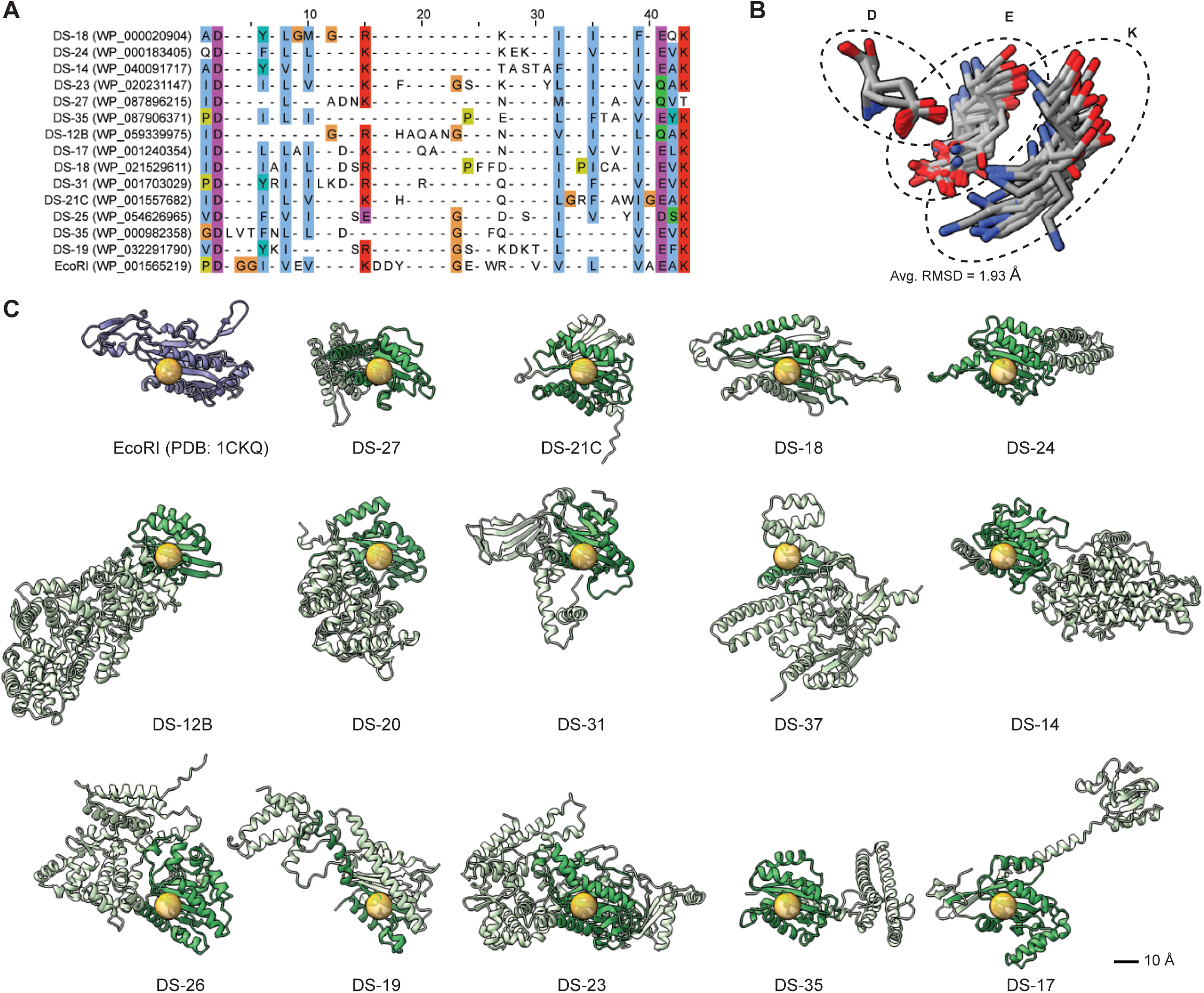
Validated PD-(D/E)XK nucleases have diverse predicted structures. (**A**) Sequence alignment of PD-(D/E)XK motifs. Protein accessions are indicated. Residues are colored based on biochemical properties using the ClustalX coloring scheme: blue – hydrophobic, red – positive charge, magenta – negative charge. (**B**) Alignment of catalytic residues from the predicted structures of the PD-(D/E)XK nucleases and the solved structure of EcoRI (PDB: 1CKQ). Aligning residues are circled and labeled with the canonical residue for that position. (**C**) Predicted structures for validated PD-(D/E)XK nucleases and the solved structure of EcoRI. The catalytic core of each protein is indicated with an orange sphere. PD-(D/E)XK domains are highlighted in green. Scale bar is shown.

**Figure S8.**
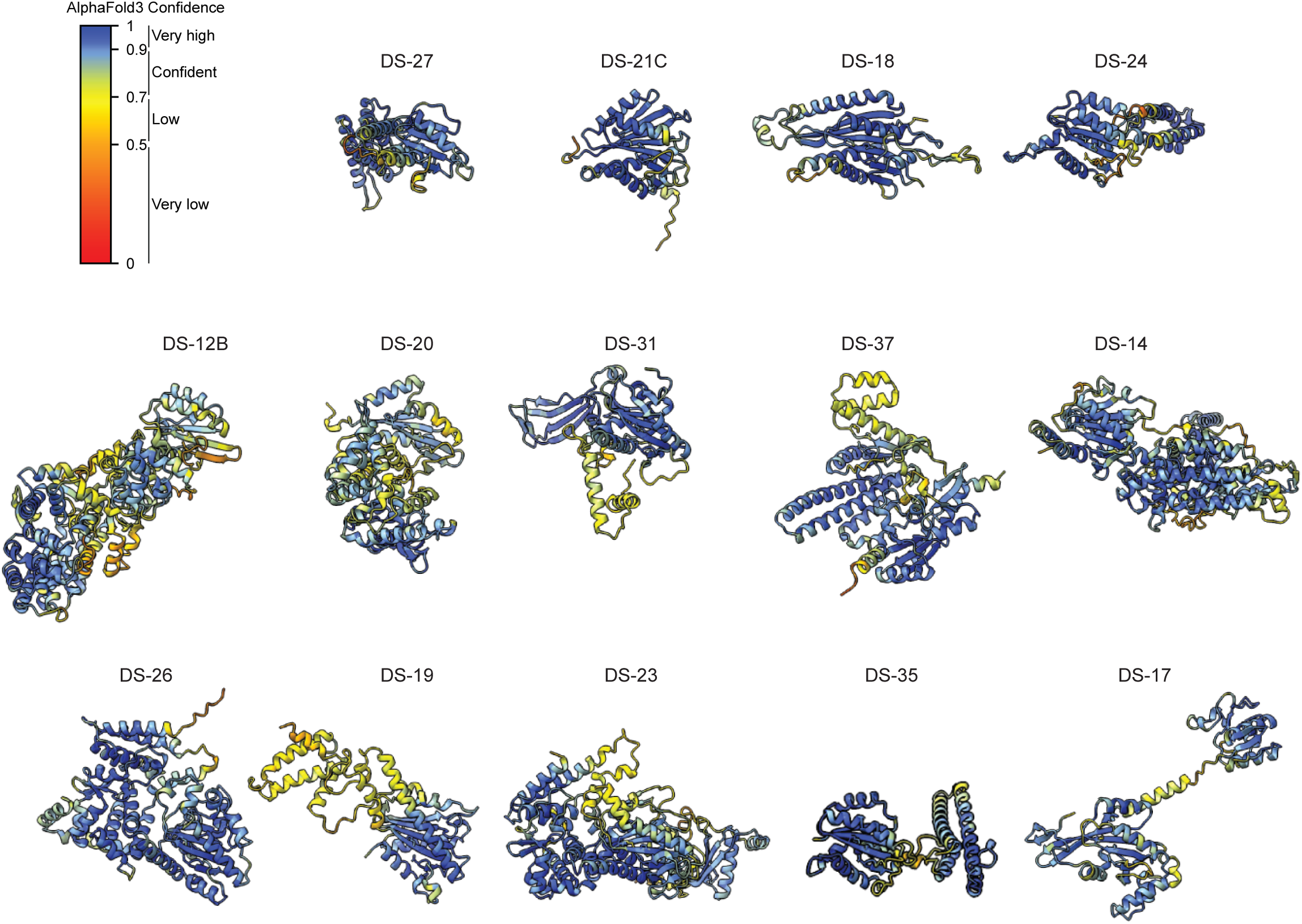
AlphaFold3 confidence metrics for PD-(D/E)XK nucleases. Per-residue pLDDT confidence scores mapped onto the predicted structures for the indicated PD-(D/E)XK nucleases.

**Figure S9.**
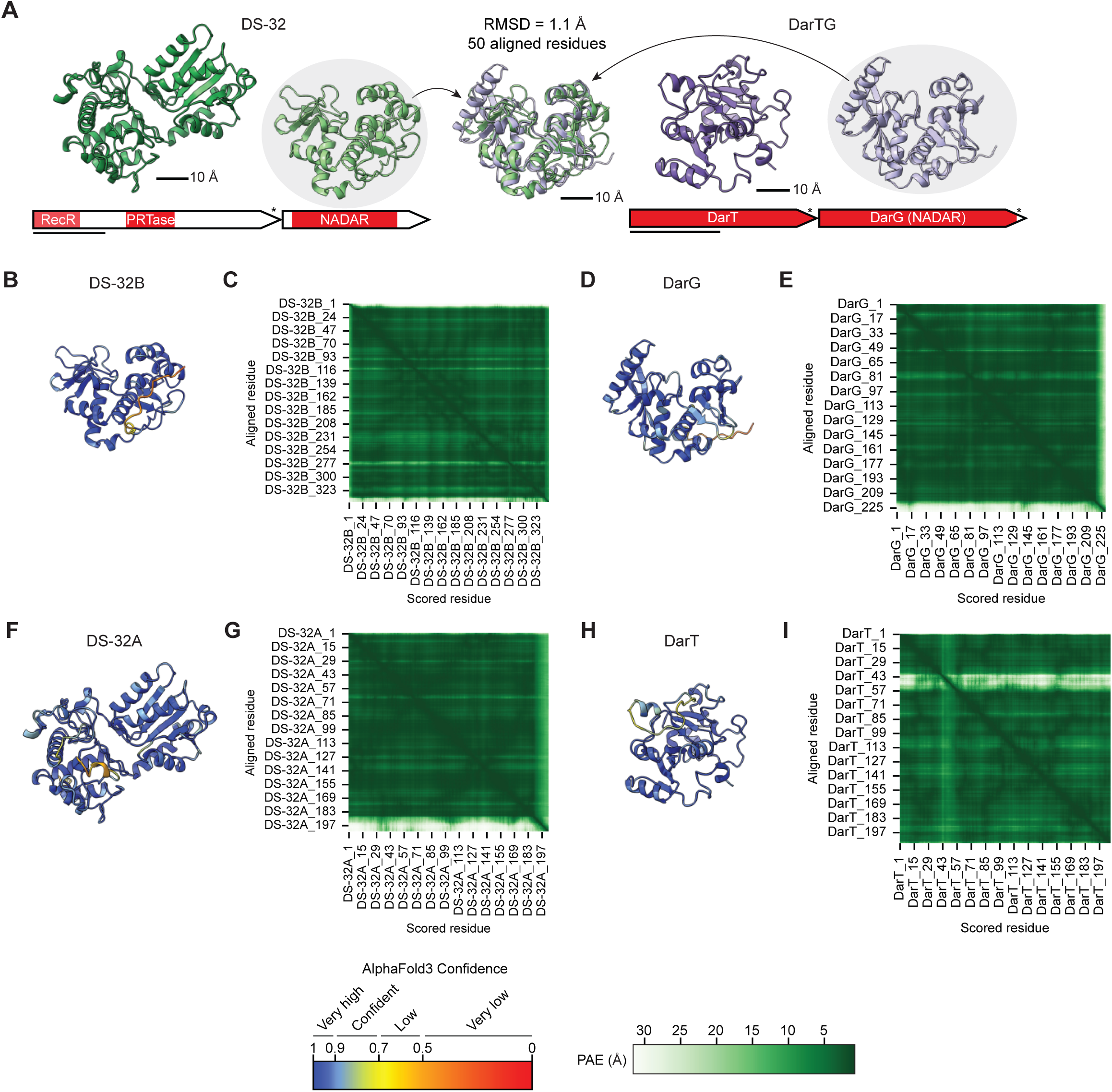
DS-32 and DarTG only have homologous NADAR proteins. (**A**) Left/bottom: domain annotations for DS-32. Left/top: predicted structures of DS-32 genes. Right: Same as left for DarTG. Center: Alignment between predicted structures of DS-32B and DarG. Scale bar is shown. (**B**, **D**, **F**, **H**) Per-residue pLDDT confidence scores mapped onto the predicted structures for DS-32B (**B**), DarG (**D**), DS-32A (**F**), and DarT (**H**). (**C**, **E**, **G**, **I**). Predicted aligned error (PAE) between residues for DS-32B (**C**), DarG (**E**), DS-32A (**G**), and DarT (**I**).

**Figure S10.**
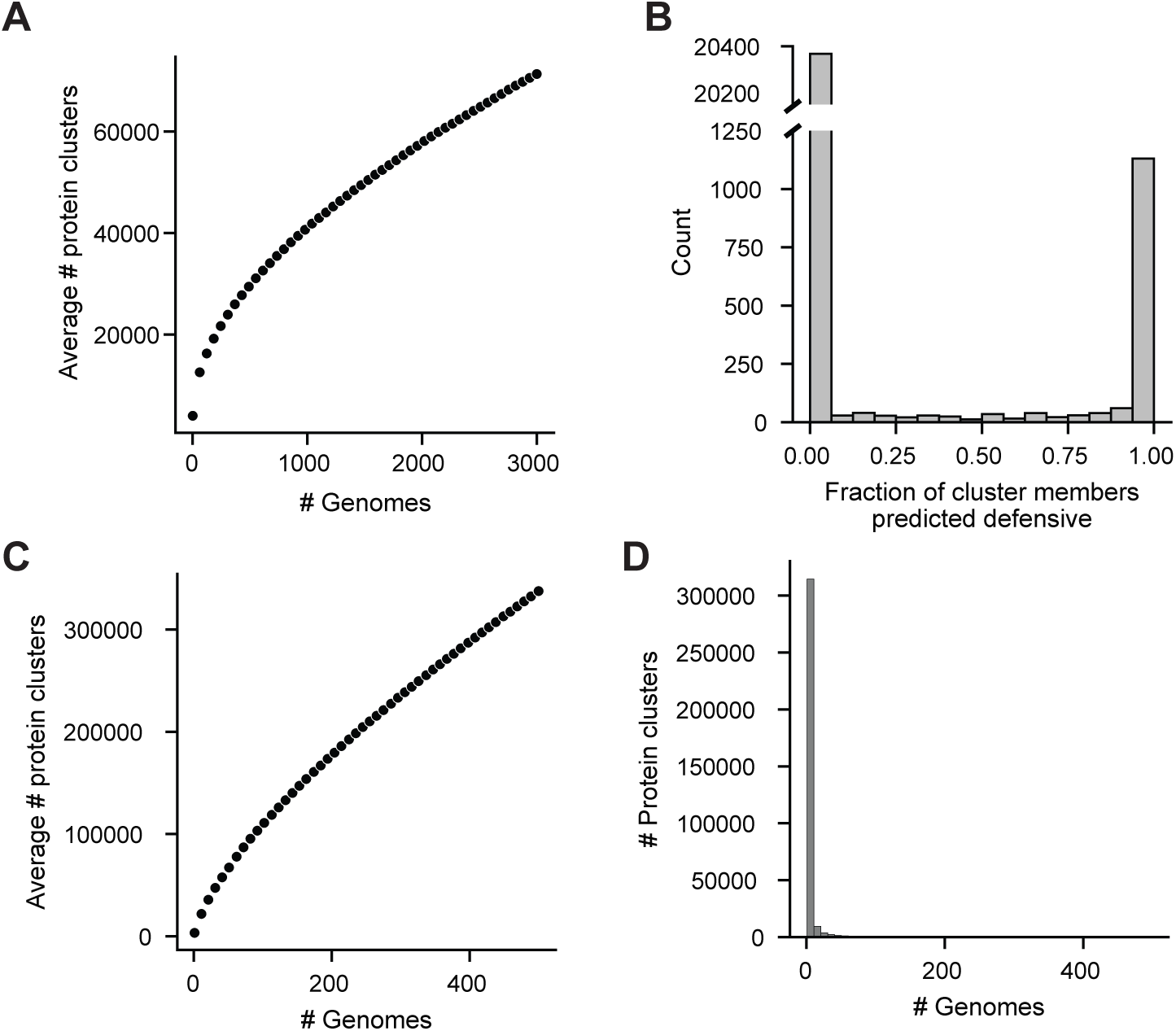
DefensePredictor outputs similar predictions for homologs. (**A**) Average number of protein clusters when considering between 1 and 3,000 *E. coli* genomes in steps of 50. Averages were calculated after resampling genomes 50 times at each step. (**B**) Histogram of the fraction of protein cluster members predicted as defensive. Only clusters with three or more members from a set of 3,000 *E. coli* strains were considered. (**C**) Same as (**A**) for 500 diverse bacterial stains. (**D**) Histogram of the number of genomes each protein cluster is encoded by in a set of 500 diverse bacterial strains.

